# Intratumoral expression of JAML on NK cells is controlled by tumor microenvironment and MHC class I interaction

**DOI:** 10.64898/2026.04.15.718645

**Authors:** Daniel Labuz, Sina Angenendt, Nora Marek, Julia Bremser, Della M. Braddish, Léa Nyman, Julian Fischbach, Ludvig Keim, Alannah Hyland, Carolina Bento, Raea Michie, Rebecca M. Lane, Lucia Carmela Passacatini, Shengduo Pei, Yueyun Pan, Mikael C. I. Karlsson, Anna Pumpe, Ann-Sophie Oppelt, Margareta Wilhelm, Chris Tibbitt, Sherwin Chan, Ulf Ribacke, Alda Saldan, Klas Kärre, Maria H. Johannson, Arnika K. Wagner, Jonathan Coquet, Benedict J. Chambers

## Abstract

Junctional adhesion molecule-like (JAML) is an adhesion molecule known to promote T cell activation and T cell-mediated tumor rejection. In the current study, we show that JAML expression is enriched on mouse intratumoral NK cells compared with splenic NK cells. JAML+ NK cells were associated with tissue residency and co-expressed the immune checkpoints PD-1 and LAG3. JAML expression could be induced on splenic NK cells by IL-2 and further enhanced by IL-21. JAML levels were inversely correlated with inhibitory signaling, as NK cells expressing self-recognizing Ly49 receptors had reduced JAML expression, suggesting regulation of JAML expression by MHC class I molecules. Interaction with the JAML ligand CXADR also reduced JAML surface expression, indicating that tumor-mediated membrane stripping may represent a mechanism of immunoediting. Although JAML RNA transcripts were detectable in human NK cells, JAML protein was found only intracellularly. Together, these findings identify the JAML-CXADR interaction as a potential regulatory pathway in NK cell-mediated killing of tumors.

## Introduction

Natural killer (NK) cells are innate lymphoid cells (ILCs) with important functions in tumor surveillance. NK cell activation is dependent on signals from activating and inhibitory receptors as well as pro-inflammatory cytokines <u>Lundqvist et al., 2026)</u>. Activating NK cell receptors can induce phosphorylation events that may result in the release of cytotoxic granules and cytokines upon interaction with ligands expressed on the tumor cells (Vivier et al., 2008). Healthy cells are protected from killing by NK cells by their expression of self-MHC class I molecules (MHC-I) on the surface which are ligands for killer-cell immunoglobulin-like receptors (KIRs) in humans, Ly49 molecules in mouse and NKG2A in both species (Hans-Gustaf Ljunggren and Klas Kärre, 1990; McQueen and Parham, 2002). Engagement of inhibitory receptors results in recruitment of phosphatases such as SHP-1, SHP-2 and SHIP-1, and dephosphorylation of signaling molecules which prevents NK cell-mediated killing (Chen et al., 2024).

For NK cells to access the TME, chemokine signals are required for recruitment, originating either from the tumor cells or from other immune populations within the tumors, such as intratumoral dendritic cells and macrophages (Wolf et al., 2023). For NK cells to exit the bloodstream, they must first adhere to the endothelial surface, subsequently slow their movement, and then transmigrate through endothelial tight junctions (Ran et al., 2022). Endothelial cells are linked to each other via intercellular junctional complexes including gap junctions, adherens junctions and tight junctions (Cong and Kong, 2020). Another group of molecules involved in migration are junctional adhesion molecules known as JAMs that are intercellular junction-associated type-I Ig superfamily proteins (IgSFs). These molecules have been shown to serve as ligands for phagocytic mononuclear (PMN) cells and monocytes as they migrate across endothelium (Ebnet et al., 2004; Marhn-Padura et al., 1998).

Junctional adhesion molecule-like protein (JAML, also known as AMICA1) is a member of the JAM family and binds to coxsackie and adenovirus receptor (CAR or CXADR) (Moog-Lutz et al., 2003; Zen et al., 2005). JAML was originally found to facilitate adhesion between tight junctions by interacting with CXADR (Luissint et al., 2008; Zen et al., 2005). However more recently, JAML expression on T cells has been shown to act as costimulatory signal and lack of JAML was associated with increased tumor outgrowth *in vivo* (McGraw et al., 2021; Witherden et al., 2010). In mouse tumor models, anti-JAML agonistic antibody therapy enhanced T cell responses against tumors, by possibly replenishing a pool of stem-like CD8+ T cells (Eschweiler et al., 2023). These data suggest that JAML might play an important role in the development of anti-tumor responses in lymphocytes.

We recently observed PD-1 expression on a distinct subset of tumor-infiltrating NK cells. Notably, these PD-1^+^ NK cells showed transcriptional clustering with the gene *Amica1* (JAML) (Wagner et al., 2022). PD-1 expression has also been associated with JAML in CD8^+^ T cells in mice (Eschweiler et al., 2023; McGraw et al., 2021). These data suggest that there is a link between JAML and PD-1 expression on intratumoral lymphocytes associated with exhaustion phenotypes. Therefore, in the current study we investigate the expression of JAML on intratumoral NK cells and which NK cell subsets express JAML. We further examine if JAML is coupled with checkpoint molecule expression on NK cells and whether JAML expression is also associated with expression of its ligand on tumors. Together, we found that JAML is expressed on a partly mature but intermediate subset of activated NK cells and disruption of the JAML-CXADR axis in NK cells drives a more immature NK cell phenotype within tumors.

## Results

### Expression of JAML on intratumoral NK cells

In our previous publication, we found an enrichment of the *Amica1*/*Jaml* transcript in a subset of tumor-infiltrating NK cells (Wagner et al., 2022), and JAML protein expression has been reported on tumor-infiltrating T cells (Eschweiler et al., 2023; McGraw et al., 2021). We therefore examined a panel of tumor cell lines to determine JAML expression on tumor infiltrating NK cells. The percentage of NK cells expressing JAML in the spleen is low which consistent with observations in T cells (Figure 1A). In mice bearing B16F10 melanoma, the proportion of tumor-infiltrating NK cells expressing JAML was slightly higher, but not significantly different from that observed in splenic NK cells, mirroring findings previously seen in T cells. Similarly, the percentage of NK cells expressing JAML in RMA-S, a lymphoma that lacks TAP2 and as such is regarded as a tumor cell line targeted by NK cells, was also not significantly different from the percentage observed on splenic NK cells (Figure 1C left panel and H). Staining of both these tumor types with JAML-Fc fusion protein revealed low to no binding, indicating a lack of detectable ligands for JAML on these cells (Figure 1 B and C right panels).

**Figure 1.**
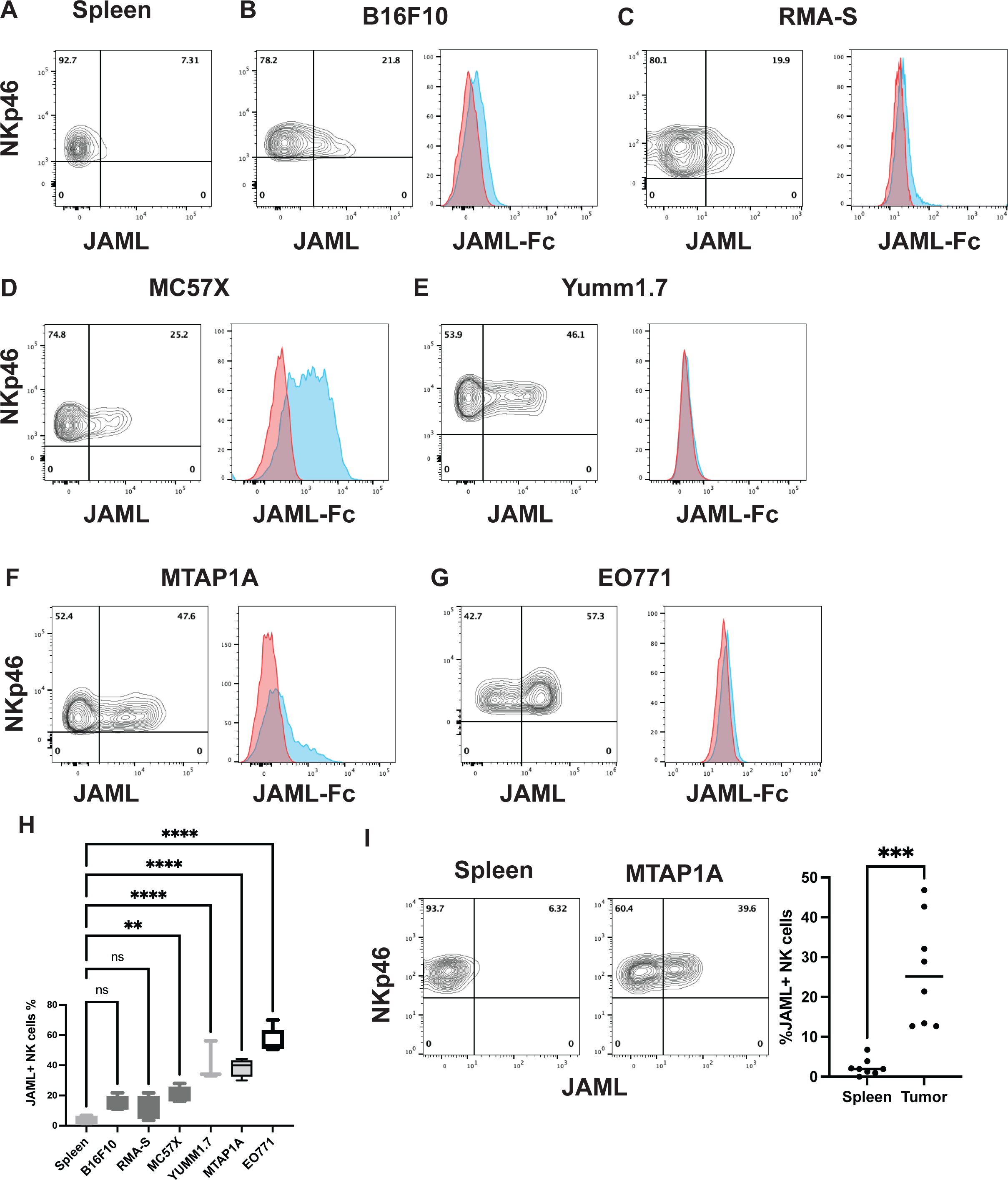
JAML expression on tumor infiltrating NK cells. (A) Expression of JAML on spleen NK cells (B) Expression of JAML on tumor infiltrating NK cells from B1610 tumors (left panel) and the binding of JAML-Fc protein to the surface of B16F10 (right panel) (C) Expression of JAML on tumor infiltrating NK cells from RMA-S tumors (left panel) and the binding of JAML-Fc protein to the surface of RMA-S (right panel). (D) Expression of JAML on tumor infiltrating NK cells from MC57X tumors (left panel) and the binding of JAML-Fc protein to the surface of MC57X (right panel). (E) Expression of JAML on tumor infiltrating NK cells from YUMM1.7 tumors (left panel) and the binding of JAML-Fc protein to the surface of YUMM1.7 (right panel). (F) Expression of JAML on tumor infiltrating NK cells from MTAP1A tumors (left panel) and the binding of JAML-Fc protein to the surface of MTAP1A (right panel). (G) Expression of JAML on tumor infiltrating NK cells from EO771 tumors (left panel) and the binding of JAML-Fc protein to the surface of EO771 (right panel). (H) Comparison of the expression of JAML on tumor infiltrating NK cells from B16F10 (n=5±SD), RMA-S (n=9), MC57X (n=5), YUMM1.7 (n=3), MTAP1A (n=9) and EO771 (n=5), statistics comparison using ANOVA comparing percentages between tumor infiltrating NK cells and splenic NK cells **p<0.001, ****p<0.0001. (I) Expression of JAML on splenic (left panel) and tumor infiltrating NK cells from MTAP1A tumors from RAG1^-/-^ mice (middle panel) and comparison in expression of JAML (right panel). ***p<.0001 Mann Whitney test (n=8). Gating strategy for NK cells can be seen in Supplemental Figure 1.

The MC57X tumor which is a methylcholanthrene induced fibrosarcoma had relatively higher levels of JAML-ligands on its surface (Figure 1D right panel). We also found that the percentage of JAML expressing NK cells within the MC57X tumors was significantly higher than that found in the spleen (Figure 1D left panel and H). When we tested another melanoma cell line YUMM1.7, the percentage of NK cells expressing JAML was significantly higher than the percentage seen on tumor infiltrating NK cells from B16F10 and splenic NK cells (Figure 1E left panel and 1H). Like B16F10 though, we found little binding of the JAML-Fc fusion protein to the surface of the tumor (Figure 1E right panel). In line with our previous scRNAseq data (Wagner et al., 2022), we detected JAML protein expression on the intratumoral NK cells from MTAP1A tumors (Figure 1F left panel and H). The MTAP1A tumor also had some binding of the JAML-Fc protein to its surface with some cells having high expression (Figure 1F right panel). Finally, tumor infiltrating NK cells from E0771 breast cancer also had a significantly higher percentage of JAML expressing NK cells than the spleens and of all our tumors tested had on average the highest percentage of JAML expressing NK cells (Figure 1G left panel and H). When stained the JAML-Fc, this tumor had low or no binding suggesting that there were no JAML ligands expressed on these tumors (Figure 1G right panel).

Since tumor infiltrating T cells could have contributed to the induction of JAML on NK cells within the tumors, we also grafted RAG1^-/-^ mice with MTAP1A. JAML expression on NK cells within the spleen were again relatively low and at levels comparable to C57Bl/6 mice (Figure1I left panel). However, the percentage of JAML expression on tumor-infiltrating NK cells was significantly higher when compared to the NK cells found in the spleen (Figure 1I right panel).

### Phenotype of JAML-expressing NK cells

We previously identified an association between JAML and PD-1 expression on NK cells (Wagner et al., 2022). Consistent with this, analysis of our scRNAseq dataset for NK cells isolated from MTAP1A tumors revealed that JAML expression was also associated with activation markers such as CD69 and the checkpoint marker LAG3, as well as tissue-resident markers such as CXCR6 and CD49a (Figure 2A). Similar findings were also observed when we examined scRNAseq dataset for tumor-infiltrating NK cells from EO771 tumors (Supplemental Figure 2A). When we performed UMAP analysis on concatenated data for NK cells from MTAP1A tumors using JAML, PD-1, LAG3, and CD69 as parameters, we observed that JAML protein expression was associated with activated CD69⁺ populations. While we did not detect any major population of TIM3-expressing NK cells, LAG3-expressing NK cells were present and were also associated with JAML expression (Figure 2). These patterns were also observed in EO771, MC57X and YUMM1.7 (Supplemental Figure 2B-D). The phenotype of the JAML⁺ NK cells demonstrated that there was a link between the expression of PD-1 and LAG3 on these cells. In general, while TIM3 expression on T cells has been associated with exhausted T cells (Das et al., 2017), we detected little or no TIM3 expression on the tumor infiltrating NK cells (Figure 2B and Supplemental Figure 2B-D).

**Figure 2.**
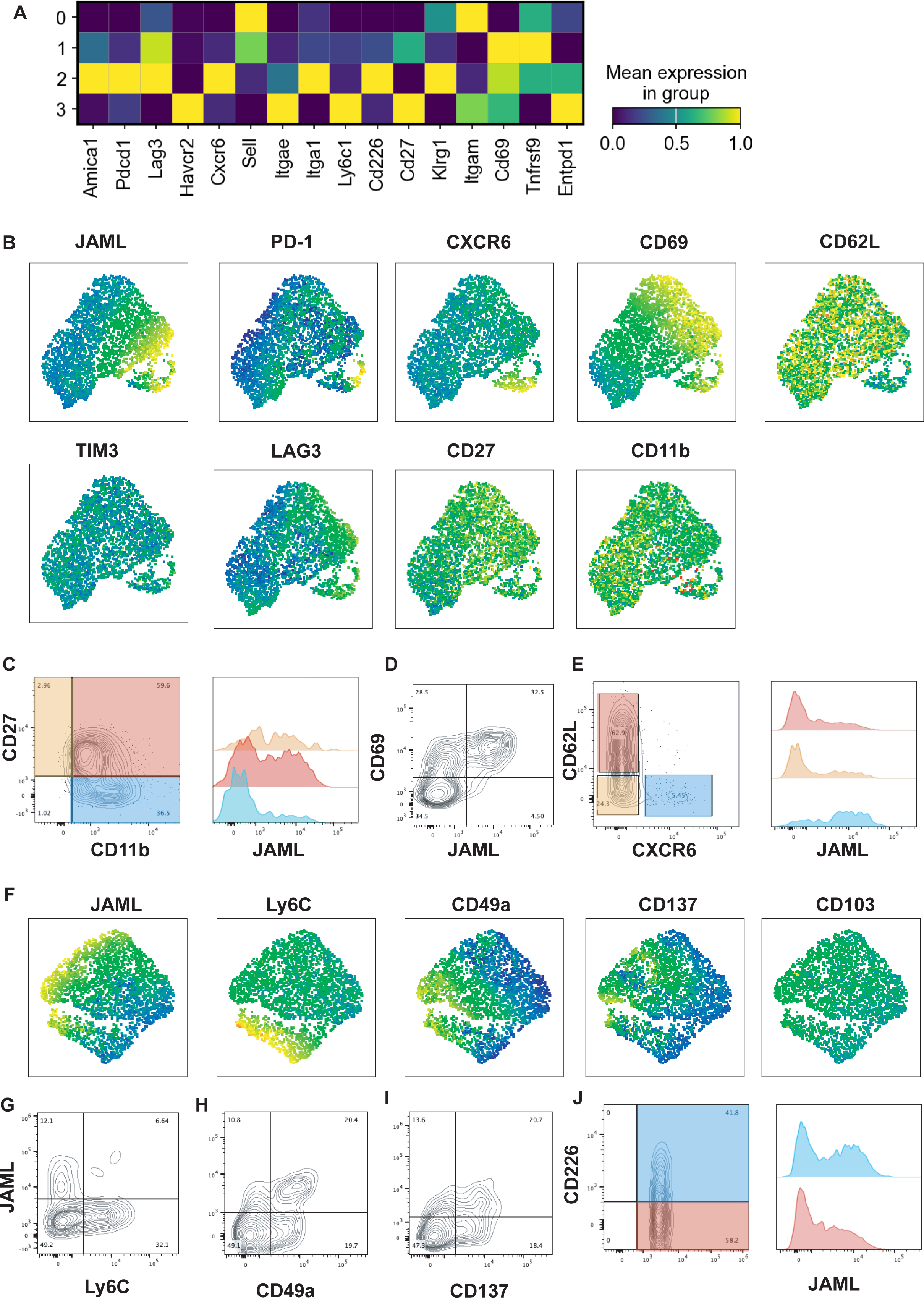
Phenotype of JAML expressing NK cells. (A) Matrix plot of scRNAseq of intratumoral NK cells from MTAP1A tumors. (B) UMAP of NK cell populations examining the distribution of JAML, PD-1, CXCR6, CD69 CD62L, TIM3, Lag3, CD27 and CD11b by flow cytometry. Data concatenated from 7 mice FACS plots showing JAML expression on concatenated data from 7 mice on (C) CD27 and CD11b cells, (D) on CD69 and (E) on CD62L and CXCR6 NK cells (F) UMAP of NK cell populations examining the distribution of JAML, Ly6C, CD49a, CD137 and CD103 by flow cytometry (G) Expression of JAML and Ly6C, (H) Expression of JAML and CD49a (I) Expression of JAML and CD137. (J) Staining of CD226 and JAML on intratumoral NK cells showing the difference in JAML expression on CD226^hi^ and ^lo^ NK cell subsets.

Maturation of NK cells can be divided into three populations CD27^+^CD11b^-^, CD27^+^CD11b^+^ and CD27^-^CD11b^+^population, where the first are more naïve and the latter are the most mature and more cytotoxic (Chiossone et al., 2009; Hayakawa and Smyth, 2006). The CD27^+^CD11b^-^ and CD27^+^CD11b^+^ NK cells had a higher frequency of JAML expression than the most mature CD27-CD11b+ population. CXCR6 has been linked to tissue residency in NK cells (Peng et al., 2013) and we have previously shown that CXCR6 expression is associated with PD-1 expression (Wagner et al., 2022). In our tumor models, CXCR6⁺ NK cells also expressed JAML (Figure 2B & E and Supplemental Figure 2). In addition, a subset of recently recruited CD62L⁺ NK cells demonstrated JAML expression (Figure 2B and Supplemental Figure 2B-D). CD49a, another marker of tissue residency, similarly co-segregated with JAML expression (Figure 2B & F). Ly6C, which has been associated with functionally exhausted NK cells (Omi et al., 2014), was also expressed by a fraction of JAML⁺ NK cells (Figure 2F & G). Interestingly, a stronger association between Ly6C and JAML was observed in EO771, MC57X and YUMM1.7 than in the MTAP1A tumors (Supplemental Figure 3A-C). Moreover, 4-1BB (CD137), which has been associated with NK cell function and favorable prognostic outcomes in tumors (Melero et al., 1998; Singh et al., 2024), identified another subset of NK cells that also expressed JAML (Figure 2F & I and Supplemental Figure 3A-C). Lastly, DNAM-1 (CD226) an activating molecule of NK cells, is preferentially expressed on JAML+ NK cells (Figure 2J).

Overall JAML^+^ NK cells were associated with tissue residency and activated cells. The low number of CD27^-^CD11b^+^ NK cells expressing JAML suggests that functionally mature NK cells do not express JAML. However, PD-1 and LAG3 were associated with JAML suggesting that these non-MHC linked inhibitory receptors on NK cells could be playing a role in controlling the NK cell activity. There were JAML^+^ Ly6C subsets and DNAM1 subsets of NK cells in the tumors suggesting that JAML was also associated with activated NK cells.

### JAML induction on NK cells *in vitro* by IL-2 and IL-21

Given the low frequency of JAML expression on splenic NK cells, we investigated whether cytokine stimulation could induce JAML on NK cells. From our previous single cell RNA-seq data, we found that the JAML positive intratumoral NK cells were associated with expression of IL-2 receptor (Wagner et al., 2022)(Figure 3A). Similarly, analysis of scRNAseq data of intratumoral NK cells from EO771 also showed that IL-2 receptors were associated with JAML (Figure 3B).

**Figure 3.**
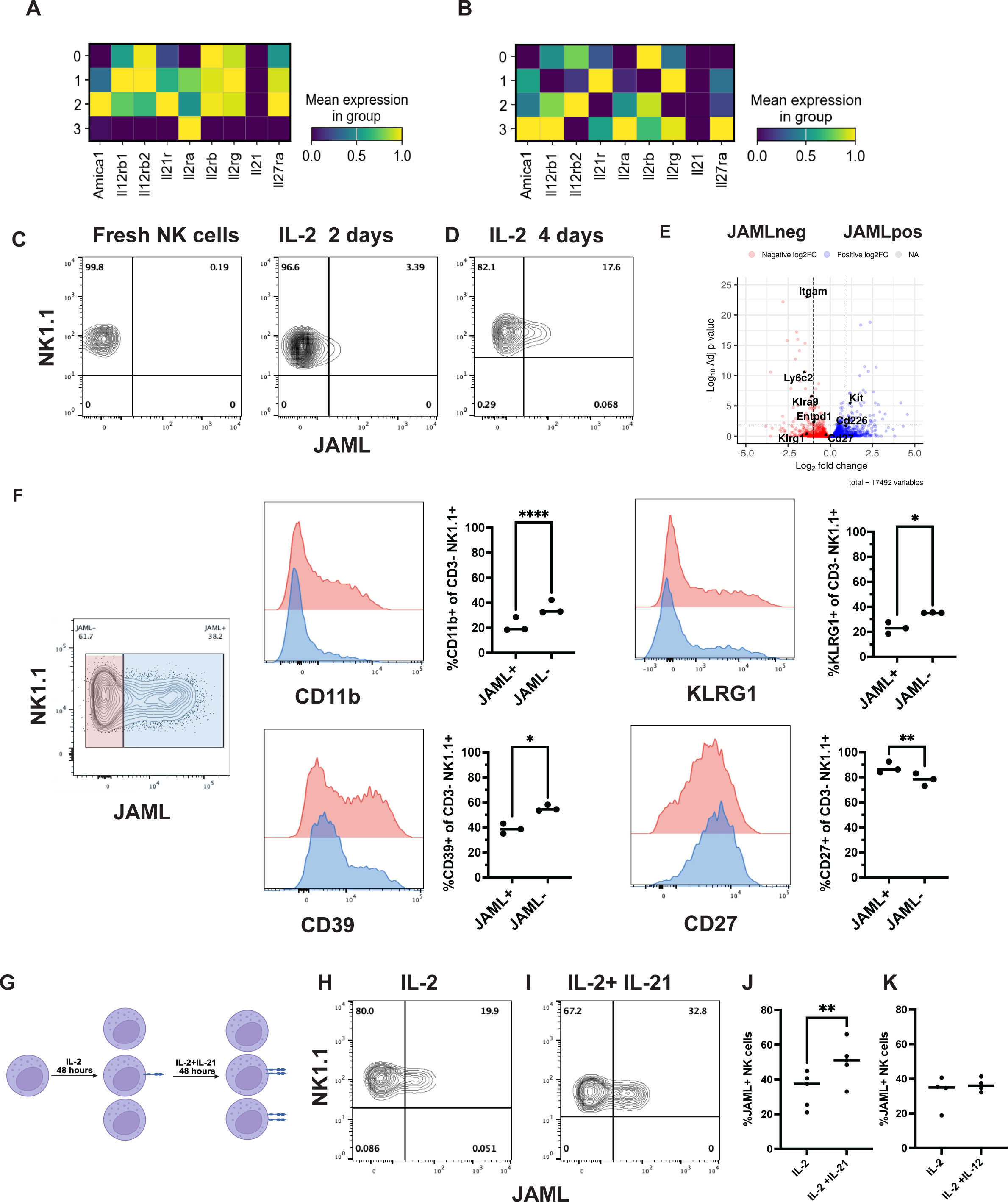
JAML is induced on NK cells by IL-2 and IL-21. Matrix plot of expression of cytokine receptors and JAML (*AMICA1*) on NK cells from (A) MTAP1A and (B) EO771. (C) Expression of JAML on fresh and IL-2 stimulated NK cells after 48 hours of culture. (D) Expression of JAML on NK cells after 72 hours of culture in IL-2. (E) NK cells cultured in IL-2 or IL-2 and IL-21 in the final 48 hours (F) JAML expression from IL-2 alone culture and (G) JAML expression from IL-2 and IL-21 culture. (H) Comparison on JAML expression between the two cultures. **p<0.01 paired t-test (n=4) separate cultures. (I) Comparison on JAML expression on NK cells cultured in IL-2 or IL-2 and IL-12 (n=3).

*In vitro* stimulation of NK cells with IL-2 resulted in JAML expression on approximately 6% of NK cells after 48 hours, with a further increase observed following an additional 48 hours of IL-2 stimulation (Figure 3C and D). When we compared the transcriptome between sorted JAML+ and JAML-NK cells, we found that CD39, KLRG1 and CD11b expression were associated with the JAML-NK cells (Figure 3E). This suggested as we observed *in vivo* that mature NK cells expressed lower levels JAML. Flow cytometric analysis confirmed lower surface expression of CD39, KLRG1, and CD11b on JAML⁺ NK cells (Figure 3F). In addition, the scRNA-seq analysis of intratumoral NK cells expressing JAML also revealed an association with the expression of the IL-21R (Figure 3A). Therefore, we also tested if IL-21 could induce JAML on NK cells. IL-21 has been shown to be a potent activator of NK cells but can also induce cell death (Brady et al., 2004), so NK cells were stimulated first in IL-2 for 48 hours and then stimulated for a further 48 hours with IL-2 again in the presence or absence of IL-21 (Figure 3G). No significant differences in cell viability were observed between the treatment conditions. In the presence of IL-21, we found on average a 33% increase in frequency of JAML expressing NK cells (33.9% JAML^+^IL-2 stimulated NK cells±10.2 vs 50.3% JAML^+^IL-2/IL-21 stimulated NK cells±11.8 p<0.01 paired t-test n=5, Figure 3H-J). In contrast, stimulation of cells with IL-2 and IL-12 had no effect on increasing JAML expression (Figure 3K).

### Control of JAML expression by inhibitory Ly49 molecules

Mouse NK cells express variable combinations of Ly49 receptors which are regulated by stochastic expression, allelic exclusion and influenced by expression of MHC class I (Brodin et al., 2009). Analysis of our scRNAseq dataset of tumor infiltrating NK cells from MTAP1A tumors revealed that cluster 1, which shows the highest *Jaml* (*Amica1*) RNA expression, showed reduced expression of the self-recognizing MHC class I inhibitory molecules Ly49C (KLRA3) and Ly49I (KLRA9) (Wagner et al., 2022) (Figure 4A). To confirm the scRNA-seq data, we performed flow cytometric analysis of intratumoral NK cells from MTAP1A tumors with a panel of Ly49-specific antibodies (inhibitory Ly49A, Ly49F, Ly49C/I (5E6 clone), Ly49I and Ly49G2 and the activating Ly49D and Ly49H) as well as NKG2A. By performing a UMAP analysis using only the inhibitory molecules on intratumoral NK cells, we observed that NK cells expressing Ly49C and Ly49I tended to have lower JAML expression and fewer cell that co-express JAML (Figure 5B). Furthermore, by comparing the staining of NK cells with the 5E6 clone that recognizes Ly49C and I with the YLI-90 clone that only recognizes Ly49I, we found that it was the double positive NK cells had a reduced frequency of JAML (Figure 4C and D). There was also significantly less expression of JAML on the Ly49C/I+ NK cells (5873±2229 MFI Ly49C/I-NK cells vs 3184±1379 MFI Ly49C/I+ NK cells *p<0.05 Wilcoxon t-test).

**Figure 4.**
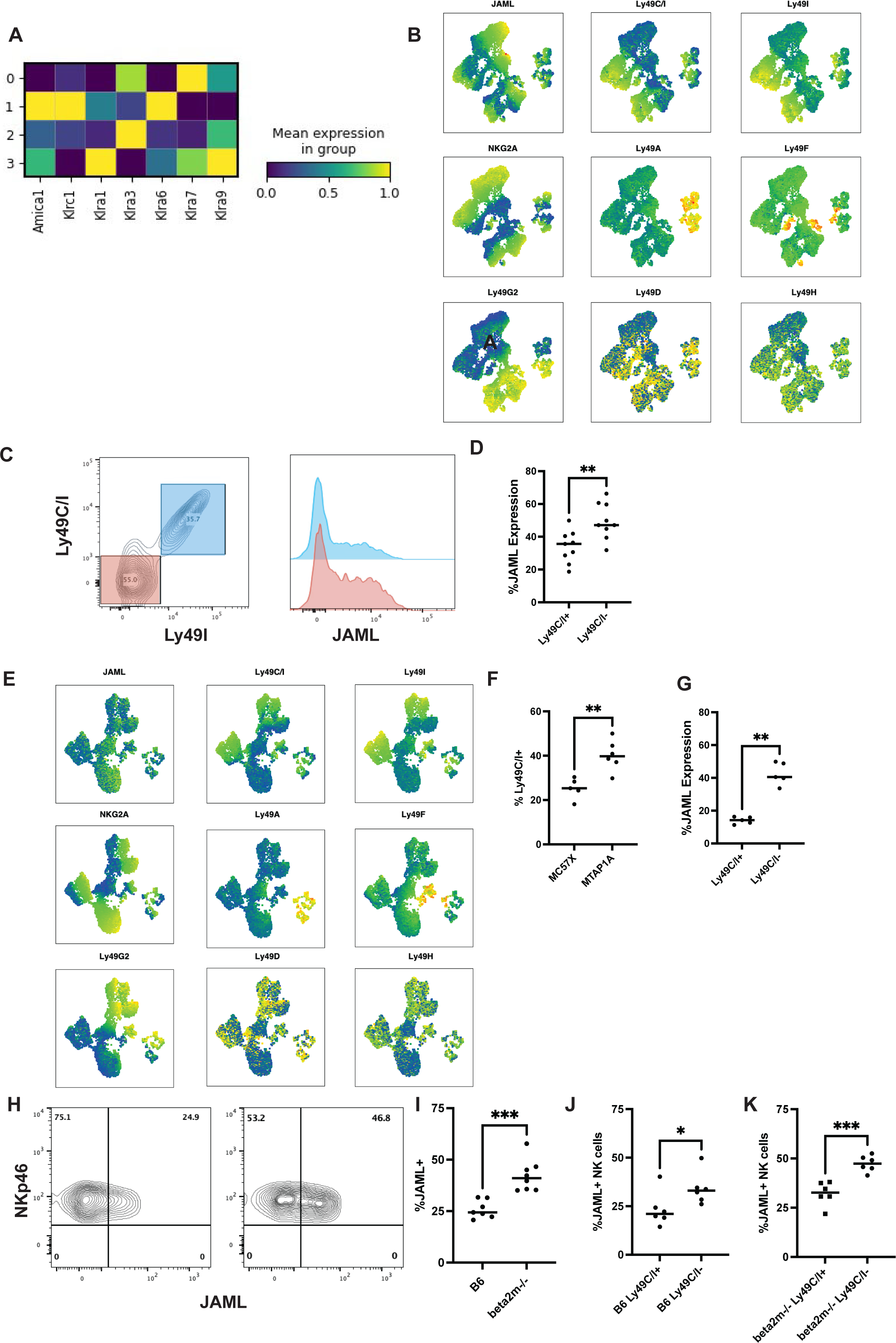
JAML expression is controlled by Ly49 expression of the NK cells. (A) Matrix plot of expression of cytokine receptors and JAML (*AMICA1*) on NK cells from MTAP1A. (B) UMAP plots of JAML, the inhibitory Ly49s Ly49C/I, Ly49I, NKG2A, Ly49A, Ly49F, Ly49G2 and the activating Ly49s Ly49D and H. UMAP was performed using the JAML and the negative Ly49s. (C) Expression of JAML on Ly49C/I NK cells. (D) Comparison of JAML expression on Ly49C/I+ NK cells vs Ly49C/I-NK cells. (**p<0.01 Mann Whitney test n=9) (E) UMAP plot of NK cells from MC57X tumors of JAML, the inhibitory Ly49s Ly49C/I, Ly49I, NKG2A, Ly49A, Ly49F, Ly49G2 and the activating Ly49s Ly49D and H. UMAP was performed using the JAML and the negative Ly49s. (F) Comparison of Ly49C/I+ NK cells between MC57X and MTAP1A (**p<0.01 Mann Whitney test n=5 and 7). (G) Comparison of JAML expression on Ly49C/I+ NK cells vs Ly49C/I-NK cells. (**p<0.01 Mann Whitney test n=5). (H) Expression of JAML on splenic NK cells isolated B6 (left panel) and β2m-/- mice stimulated with IL-2. (I) Comparison of JAML expression of splenic NK cells isolated B6 and β2m-/- mice (**p<0.01 Mann Whitney test n=8). (J) Comparison of JAML expression on Ly49C/I+ IL-2 stimulated NK cells vs Ly49C/I- IL-2 stimulated NK cells. (**p<0.01 paired t-test n=6) (K) Comparison of JAML expression on Ly49C/I+ IL-2 stimulated NK cells vs Ly49C/I-IL-2 stimulated β2m-/- NK cells. (***p<0.001 paired t-test n=6).

**Figure 5.**
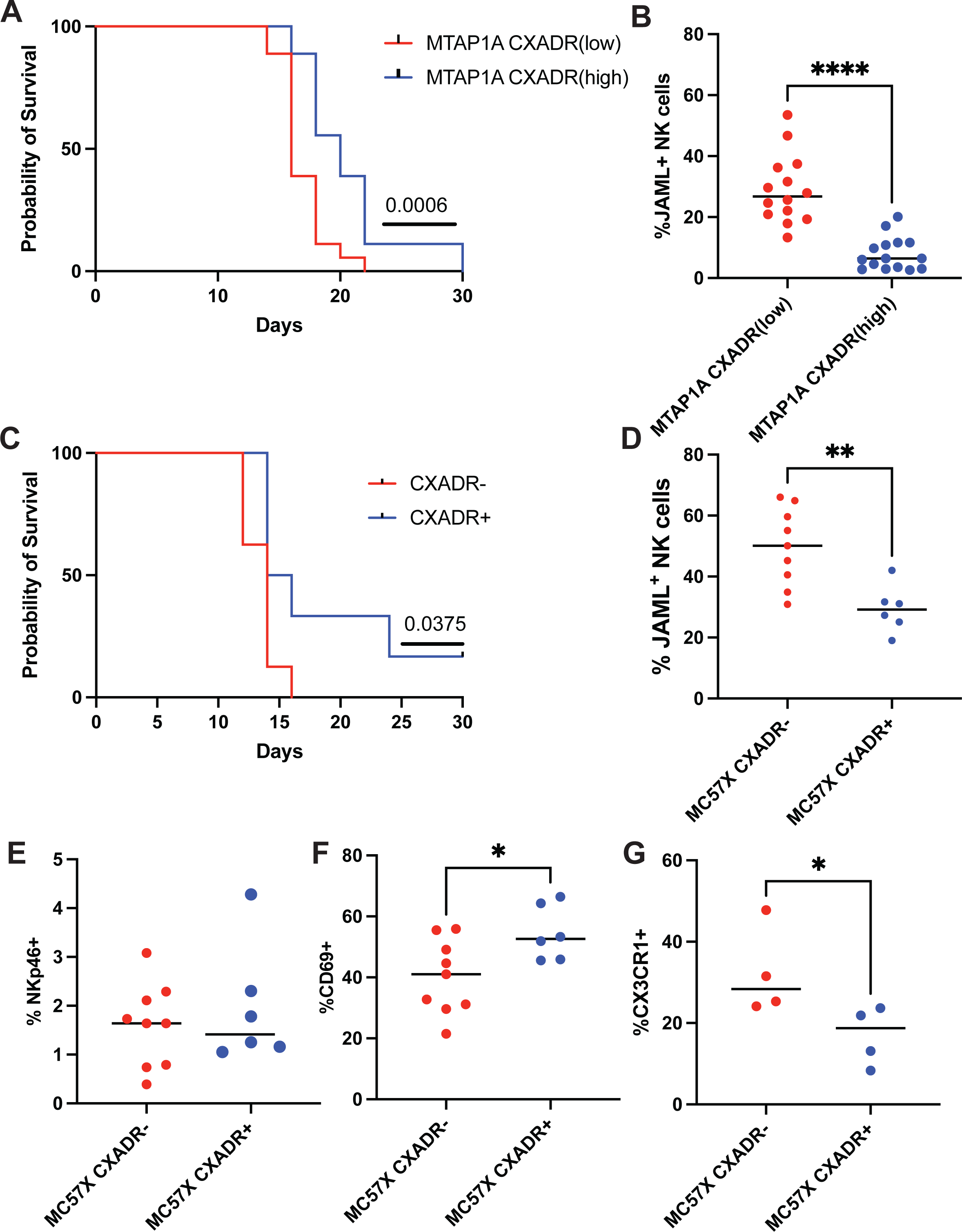
CXADR expression on tumors increases survival while reducing JAML expression on NK cells. (A) MTAP1A tumor cells were sorted into high and low expressing cells and grafted s.c. at 10^5^. Survival was increased in mice that received tumors expressing high levels of CXADR. (p<0.001 Logrank (Mantel-Cox) test n=18 mice). (B) Expression of JAML on NK cells was reduced on NK cells recovered from the MTAP1A tumors expressing high levels of CXADR. (C) MC57X tumor with CXADR knocked out had reduced survival compared to MC57X with high CXADR expression (p<0.05 Logrank (Mantel-Cox) test 14-15 mice). (D) Expression of JAML on NK cells isolated from MC57X^CXADR-^ and MC57X^CXADR+^ tumors. (E) Comparison of % NK cells in MC57X^CXADR-^and MC57X^CXADR+^ tumors, (F) CD69^+^ and (G) CX3CR1^+^ NK cells (*p<0.05 Mann-Whitney test, **p<0.01 Mann Whitney test).

Since MTAP1A expresses low surface levels of MHC class I, we next examined JAML expression on NK cells in MC57X tumor which has high surface levels of MHC class I (Figure 4E). Like the MTAP1A intratumoral NK cells, we found that Ly49C and I positive NK cells had reduced expression of JAML. Interestingly in these tumors with high MHC class I expression, there was also reduced percentage of Ly49C/I expressing NK cells compared to the tumor MTAP1A with low MHC Class I expression (Figure 4F). Furthermore, as seen in MTAP1A, the LY49C/I+ NK cells show a clear reduction in JAML expression compared to the LY49C/I-cells (Figure 4G).

To assess if this observation extends beyond fibrosarcoma tumors, we analyzed intratumoral NK cells from EO771 breast cancer. ScRNA-seq analysis suggested mutual exclusivity between JAML expression and Ly49C/I expression (Supplement Figure 4A). We confirmed this finding using flow cytometry and saw that the Ly49C/I positive NK cells had reduced expression of JAML (Supplemental Figure 4B)

To further investigate if interaction of Ly49 receptors with MHC class I molecules could control JAML expression, we stimulated wild-type NK cells and NK cells from β_2_m-deficient mice, which lack surface expression of both classical and non-classical MHC class I molecules, with IL-2 and assessed their JAML expression. We found that there was a higher percentage of JAML expressing NK cells in the β_2_m-deficient NK cells stimulated with IL-2 compared to wild-type NK cells (Figure 4H and I). This suggests that in the absence of either stable classical or non-classical MHC class I molecules, and the resulting weakened inhibitory signaling in NK cells, leads to increased JAML expression. Similar to the findings *in vivo*, we further found that JAML-expressing NK cells were reduced in populations where Ly49C/I was co-expressed on the NK cells (Figure 3I). The same observation was also made in the β_2_m-deficient mice, suggesting this might be an intrinsic effect in NK cells that express receptors for self-MHC class I (Figure 4I).

### Loss of CXADR expression on tumors leads to reduced survival and alters NK cell phenotype

Since MTAP1A cell exhibit a heterogenous surface expression of CXADR (Figure 1F), we sorted MTAP1A into high and low expressing CXADR MTAP1A cells and grafted them to mice. Mice given MTAP1A tumors with high expression of CXADR (MTAP1A^CXADRhigh^) survived significantly longer and had a slower tumor outgrowth than those mice that received the MTAP1A with low expression of CXADR (MTAP1A ^CXADRlow^) (Figure 5A and Supplemental Figure 5A)). In addition, mice that were grafted with MTAP1A^CXADRhigh^ had on average lower frequency of JAML expressing NK cells (Figure 5B).

These data suggested a role for CXADR in survival and in addition that CXADR affects JAML levels on NK cells. We generated CXADR-knockout (KO) MC57X tumor cells using CRISPR/Cas9 and grafted these CXADR-WT or CXADR-KO cells into mice. Mice receiving CXADR-KO MC57X tumor cells (MC57X^CXADR-^) displayed significantly reduced survival and accelerated tumor growth compared with those engrafted with CXADR-WT MC57X tumors (MC57X^CXADR+^) (Figure 5C and Supplemental Figure 5B). We further found that the NK cells from the MC57X^CXADR+^ tumors also exhibited reduced frequency of JAML expressing NK cells compared to the NK cells from the MC57X^CXADR-^ tumors (Figure 5D). In addition, while there was no difference in the frequency of tumor infiltrating NK cells between MC57X^CXADR^ and MC57X^CXADR+^(Figure 5E), MC57X^CXADR-^ tumors had lower frequency of CD69^+^ NK cells but higher frequency of CX_3_CR1^+^ on NK cells from these tumors (Figure 5F and G). This suggested that engagement of CXADR could lead to different NK cells activation state, maturation, and effector function.

### Loss of JAML expression on NK cells in the presence of CXADR is due to membrane shedding

Given the reduced JAML expression on NK cells from tumors expressing CXADR, we examined potential mechanisms underlying this effect. Initially we compared the expression of JAML on IL-2 stimulated NK cells following short-term incubation with either B16F10 tumor cells, which lack detectable JAML ligands, or MTAP1A CXADR^high^ tumor cells (Figure 6A). We found that within three hours of co-culture with MTAP1A but not B16F10, the expression of JAML had been significantly reduced on the NK cells (Figure 6B). Notably, this reduction was reversible, when these NK cells were re-cultured for 48 hours in IL-2, the percentage of JAML expressing NK cells recovered (Figure 6C). Furthermore, when IL-2 stimulated NK cells were mixed with CXADR^+^ or CXADR^-^ MC57X cells, NK cells incubated with CXADR^+^ MC57X had decreased JAML^+^ frequency compared to NK cells incubated with CXADR^-^MC57X (Figure 6D).

**Figure 6.**
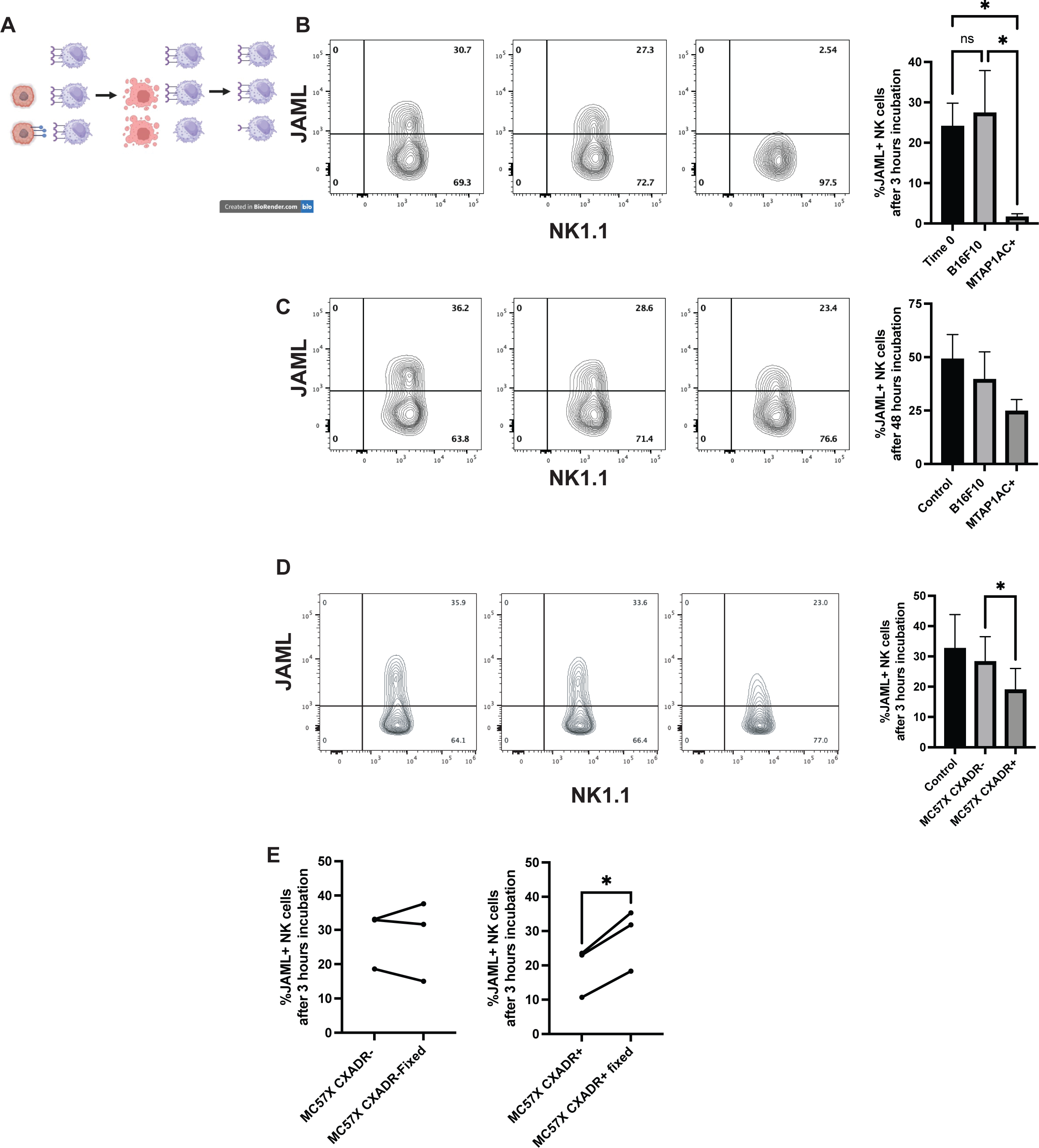
JAML is shed from the surface of NK cells upon interaction with its ligands. (A) NK cells were cocultured with tumors for three hours and then removed for staining or re-culture in IL-2. (B) Expression of JAML on IL-2 stimulated NK cells following culture in media alone, B16F10 and MTAP1A^CXADRhi^, Bar graph is representative of three experiments (*p<0.05 one way ANOVA). (C) JAML levels recover when re-cultured for 48 hours in IL2. (D) CXADR on MC57X tumors also reduce expression of JAML on IL-2 stimulated NK cells (*p<0.05 one way ANOVA) (E) Fixing tumor cells before co-incubation with tumors. Loss of JAML in the presence of fixed MC57X is ablated. (p<0.05 paired t-test n=3 experiments).

Membrane shedding is a common upon interaction between cells and can be manipulated by tumors to evade detection (Chao et al., 2025; Kim et al., 2025). We therefore evaluated whether proteolytic cleavage contributed to the loss of JAML. However, treatment with the metalloprotease inhibitor TAPI-2, which had previously been shown to prevent JAML cleavage on the surface of neutrophils (Zen et al., 2005), did not prevent loss of JAML expression on NK cells co-cultured with CXDAR^+^ MC57X (Supplementary Figure 6A). This suggested that the reduction of JAML on the surface might not be due to surface cleavage by proteases. Next, we tested if trogocytosis might be involved. Trogocytosis is a process in which immune cells physically acquire fragments of membrane from cells they contact, a process whereby NK cells remove ligands such as MHC class I or costimulatory molecules from tumor cells during immune synapse formation. We first coated plates with CXADR fusion protein and incubated NK cells for three hours after which we saw approximately 10% fewer JAML expressing NK cells (Supplementary Figure 6B). However, when we prefixed the tumor cells in formaldehyde, we failed to see reduction in the frequency of JAML expressing NK cells (Figure 6E). These data suggested that membrane interaction between the NK cells and tumors was important for JAML loss, but that loss could occur even upon contact with recombinant CXADR. Loss of JAML on the membrane of NK cells *in vivo* and *in vitro* suggest that the reduced frequency of JAML-positive NK cells in tumors might be due to the cytokine microenvironment and interaction between JAML and its ligand within the tumors.

### CXADR triggers NK cell mediated killing *in vitro*

Since mice that received tumors expressing CXADR showed slower growth than mice that receiving CXADR^low^ or ^negative^ tumors, we checked *in vitro* whether NK cells could differentiate between these tumors in a killing assay. IL-2 stimulated NK cells were mixed with MC57X and MTAP1A tumors expressing or not expressing CXADR. On average, 39% of NK cells used for the cytotoxicity assay expressed JAML on their cell surface. From four experiments, we found that on average MC57X^CXADR+^ were more sensitive to NK cells than MC57X^CXADR-^. At the 10:1 ratio, the killing was 28.7±19.2 for the MC57X^CXADR-^ compared with 53.9±9.9 MC57X^CXADR+^ (Figure 7A). Across all E:T ratios, MC57X^CXADR+^ was more sensitive to NK cells (Figure 7B). Unsurprisingly, the MTAP1A tumors due to its lack of MHC class I were more sensitive to NK cell mediated killing. However, even with these tumors at the 10:1 ratio, the killing was 64.4±16.7 for the MTAP1A^CXADR-^ compared with 74.1±15 MTAP1A^CXADR+^ (Figure 7C). Across all E:T ratios on average the MTAP1A^CXADR+^ were more sensitive to NK cells (Figure 7D).

**Figure 7.**
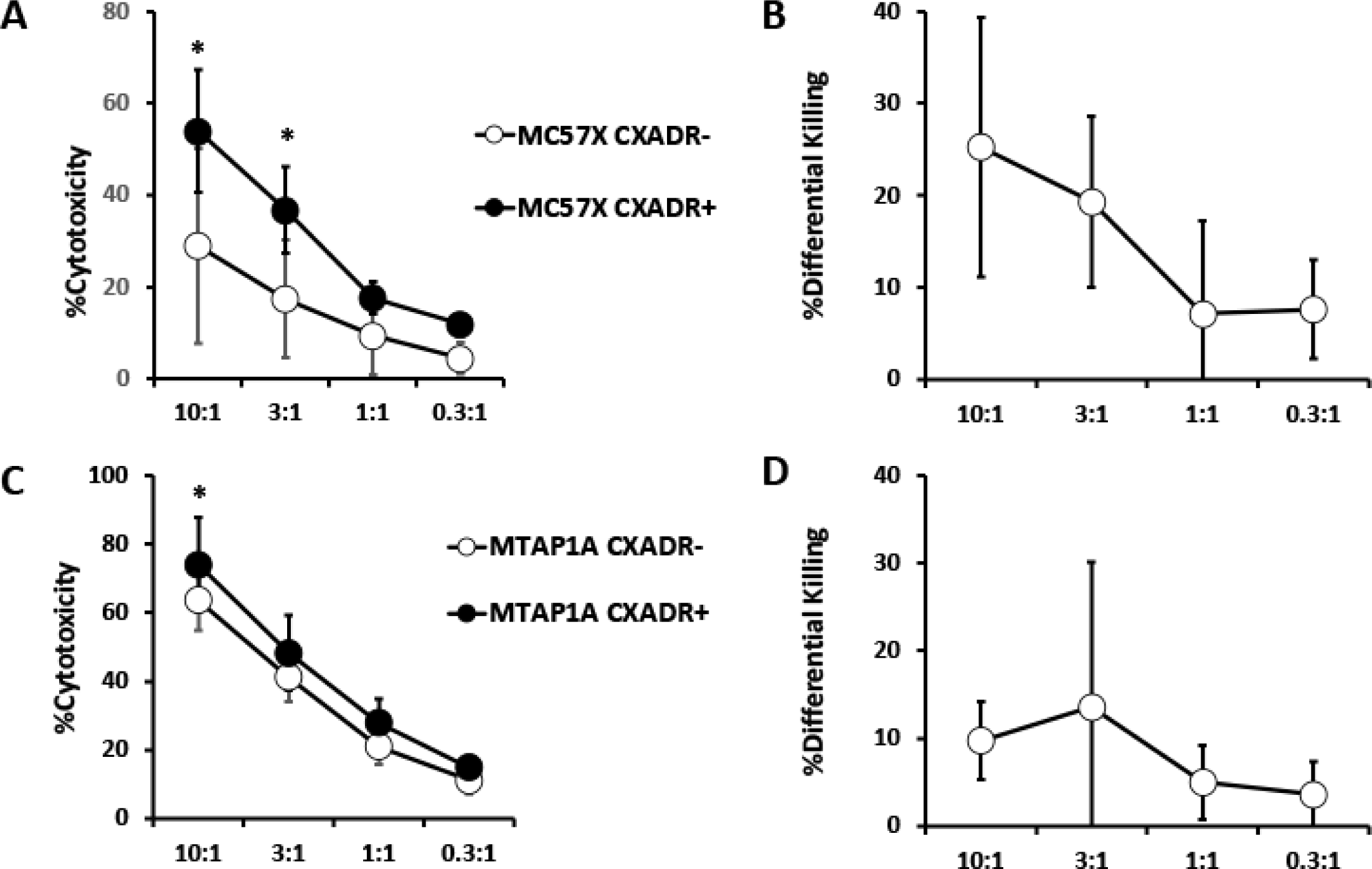
CXADR expression on MC57X and MTAP1A enhances killing of tumors by NK cells. (A) Cytotoxicity curves for IL-2 stimulated NK cell mediated killing of MC57X^CXADR+^ (*p<0.05 paired t-test N=4 separate experiments) (B) Differential killing of the MC57X^CXADR+^ compared to the MC57X^CXADR^. (C) Cytotoxicity curves for IL-2 stimulated NK cell mediated killing of MTAP1A^CXADR+^ *p<0.05 paired t-test N=4 separate experiments) (D) Differential killing of the MTAP1A^CXADR+^ compared to the MTAP1A^CXADR-^. p<0.05 paired t-test, n=4 experiments.

### JAML is expressed on several subsets of human tumor infiltrating NK cells

Previously *JAML* transcript has been shown to be enriched on intratumoral human CD8^+^ tissue resident memory T cells (Eschweiler et al., 2023). To validate and extend our observations in tumor-infiltrating NK cells in mouse tumors to human NK cells, we analyzed a pan-cancer NK cell dataset spanning seven solid tumor types from 427 patients to characterize *JAML* expression and the phenotype of *JAML*⁺ NK cells. (Netskar et al., 2024). Across tumor types, *JAML* expression was low in NK cells (∼5-10%), with no individual tumor having a significantly higher proportion of *JAML* expressing NK cells, although melanoma and sarcoma show comparatively higher expression (Figure 8A). UMAP and Leiden clustering of the tumor-infiltrating NK cells show 7 clusters, none of which was strongly enriched for *JAML* expression (Figure 8B). Next, we asked whether tumor-infiltrating JAML-expressing NK cells also expressed markers associated with tissue residency such as *CXCR6* and *ITGAE*. Cluster 3, which showed the highest relative JAML expression, showed higher levels of *ITGAE* and *ITGA1* compared to the other clusters (Figure 8C). Next, we subset the JAML^+^ and JAML^lo/-^ NK cells from tumor tissue based on an expression of 0.1 or higher (determined from violin plots across all tumors, JAML^+^ = 1234 cells and JAML^lo/-^ = 38081 cells) and saw that JAML expressing tumor infiltrating NK cells expressed higher levels of *PDCD1* and *CD27* and lower levels of *ITGAM* like what we have previously shown in mouse tumors (Figure D). Using the same dataset, we looked at healthy tissue NK cells and their levels of *JAML* and used UMAP and Leiden clustering to generate 6 NK cell clusters with cluster 3 being enriched for *JAML* compared to the other clusters and contained mostly in the prostate tissue (Figure E). We further looked at this cluster and saw that it showed higher levels of *CXCR6, CD69, CD226, CD27, and SELL,* suggesting a heterogenous NK cell population with characteristics of both tissue residency and circulating NK cells (Figure 8F). Due to the low abundance of *JAML* transcript in NK cells in human scRNAseq data, we further investigated the isoform transcript levels across several sorted immune cell subsets from healthy blood (Figure 8G) (Monaco et al., 2019). Examining the top 8 expressing *JAML* isoforms, human NK cells express very little transcript of these *JAML* isoforms, meanwhile myeloid subsets express JAML at high levels, suggesting that this protein is not likely to be expressed under homeostatic conditions (Figure 8G). To confirm this, we stimulated healthy human peripheral blood NK cells for 48 hours with IL-2 and saw no surface JAML expression but did find JAML expression within the cell (Figure 8H). Imaging flow cytometry following IL-2 stimulation for 48 hours further confirmed intracellular localization of JAML with heterogenous expression levels across the NK cells, suggesting that this protein may be subject to intracellular retention, cleavage or impaired trafficking to the cell surface (Figure 8I).

**Figure 8.**
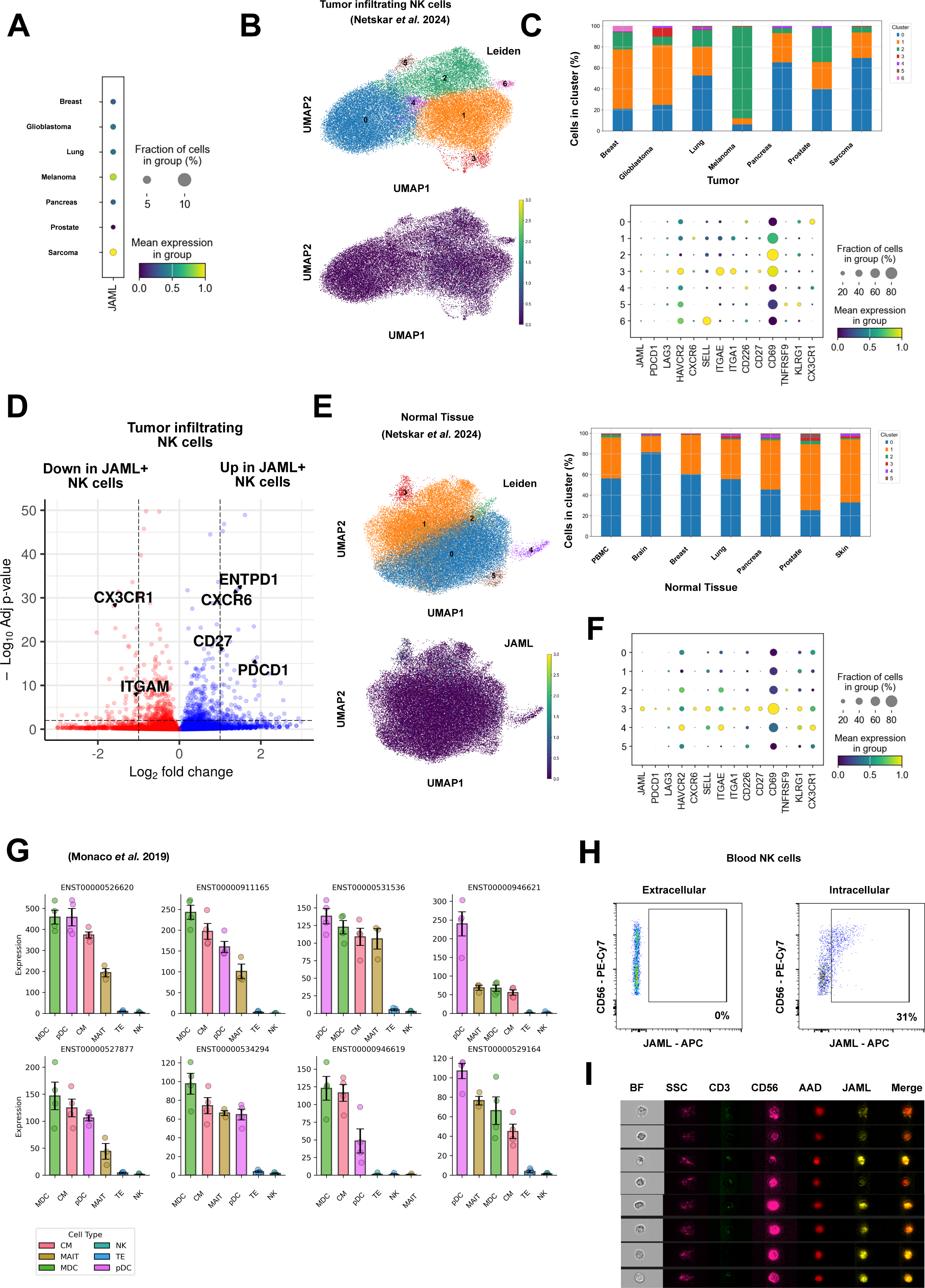
A small population of intermediate mature tumor infiltrating Human NK cells express JAML. (A) A pan-cancer scRNA-seq dataset of seven tumors was analyzed for the expression of *JAML* on tumor infiltrating NK cells. (B) UMAP defining seven clusters of the tumor infiltrating NK cells and the expression of *JAML* across the clusters with bar plots showing the contribution of each cluster in each tissue. (C) Dotplot showing genes correlated with NK cell maturation, exhaustion, activation, and tissue residency in the seven clusters of tumor infiltrating NK cells. (D) Volcano plot of a pseudobulk differentially expressed genes between JAML+ and JAML-tumor infiltrating NK cells. (E) UMAP defining six clusters of the NK cells in healthy tissue and the expression of *JAML* across the clusters with bar plots showing the contribution of each cluster in each tissue. (F) Dotplot showing genes correlated with NK cell maturation, exhaustion, activation, and tissue residency in the six clusters of healthy tissue NK cells. (G) Bar plots of selected immune cells (MDC – Myeloid dendritic cells (CD11c+ HLA-DR+), CM – Classical monocytes (CD14+ CD16-), pDC – plasmacytoid dendritic cells (CD123+ HLA-DR+), MAIT - Mucosal associated invariant T cell (Vα7.2, CD161^hi^), TE - Terminal Effector CD8+ (CCR7-CD45RA+), NK - Natural Killer Cells (CD56+ CD16+) and their expression (in transcripts per million) of the top 8 expressing *JAML* isoforms. (H) FACS plots showing the extracellular and intracellular expression of JAML in NK cells from healthy human blood following 48 hours of IL-2 stimulation. (I) Imagestream pictures showing the location of the binding of intracellular anti-JAML within healthy human blood NK cells following 48 hours of IL-2 stimulation.

## Discussion

In the present study we demonstrate that JAML expression in NK cells is closely associated with tumor infiltration. While only a small subset of splenic NK cells expressed JAML, the percentage of JAML-expressing intratumoral NK cells was generally higher, with the exception of B16F10 and RMA-S tumors. Among the tumors analyzed, EO771 had the highest frequency of JAML-expressing NK cells followed by MATP1A, MC57X and YUMM1.7. Notably, JAML frequency increased on tumor-infiltrating NK cells when CXADR was knocked out in MC57X. This pattern of expression was similar to that observed with T cells where JAML expression was also associated with tumors lacking CXADR (McGraw et al., 2021). These findings suggest that JAML expression could be controlled by the availability of its ligand on the tumor cells, although the lack of JAML induction on NK cells in B16F10 and RMA-S indicates that additional factors such as the intratumoral environment might also play a role in triggering JAML expression. Indeed, our *in vitro* data show that IL-2 and IL-21 can drive JAML expression on NK cells *in vitro*, suggesting the cytokine environment within the tumors might also be an important regulatory component of JAML expression on NK cells. Furthermore, self MHC-binding inhibitory receptors also could control JAML expression on NK cells, underscoring the role of the environment within the tumors in shaping JAML expression.

While the frequency of JAML-expressing NK cell between tumors varied, these JAML-expressing NK cells did share phenotypes in common. JAML-expressing NK cells were in general CD27^+^CD11b^-^ or CD27^+^CD11b^+^ suggesting that JAML levels were reduced on mature NK cells. Furthermore, checkpoint molecules such as PD-1 and LAG3 were also associated with the JAML population. PD-1 expression and JAML have previously been associated with T cells, however only a small subset of NK cells expresses PD-1. Within our studies, the EO771 tumor, which had the highest frequency of JAML, also had the highest frequency of PD-1 on tumor-infiltrating NK cells. However, the fact that PD-1 and JAML expression are correlated suggests that PD-1 expression on NK cells could be related to post CXADR-JAML or lack of CXADR-JAML interactions by the NK cells. Thus, NK cells express PD-1 on their surface particularly if CXADR is lost on the tumor cells and the levels of JAML on the NK cells are maintained to dampen the NK cell activity. Furthermore, we also found subtle differences in the NK cells from CXADR^+^ and CXADR^-^ tumors. Tumor-infiltrating NK cells from CXADR^+^ tumors expressed more CD69 and less CX_3_CR1, a receptor that is expressed on a later stage of differentiated NK cells in healthy tissues (Ponzeva et al., 2013), suggesting that not only can CXADR expression be driving tumorigenesis but can also affect the maturity of the NK cells within the tumor.

Inhibitory receptors on NK cells play key roles in development and maturation of NK cells. We found that within the different tumors, that Ly49 expression patterns varied in a manner consistent with the expression levels of the MHC class I molecules on the surface of the tumors. The expression of the self-recognizing Ly49 molecules Ly49C and I appeared to control expression of JAML, as cells expressing these molecules had a lower frequency of JAML expressing NK cells. NKG2A expressing NK cells had the highest expression of JAML. These findings could also be observed in IL-2 stimulated NK cells where recognition of self MHC-I impacted JAML expression. Therefore, even though JAML was induced in tumors or by cytokines, there was still an intrinsic control of its expression by MHC-I engagement.

CXADR has been shown to play a role in tumorigenesis and CXADR may enhance the invasive behaviour of cancer cells by promoting epithelial–mesenchymal transition (EMT) (Owczarek et al., 2023). In our study, CXADR expression on tumors varied markedly from tumor to tumor. Loss of CXADR has been associated with metastasis which might explain why some of our tumors in the study are CXADR negative. However, lack of CXADR on tumors such as YUMM1.7, E0771, RMA-S and B16F10 might also suggest a selection pressure since these tumors have been used extensively for *in vivo* tumor studies. Our two methylcholanthrene induced tumors MC57X and MTAP1A had a consistent expression CXADR even if MTAP1A could be divided into high and low expressers. Since these fibrosarcoma tumors were induced in the skin, it might explain their expression of CXADR compared with the other tumors that we examined. Moreover, both MTAP1A and MC57X have not been used extensively compared to B16F10 and E0771 and so may not have undergone selection to lose CXADR. Indeed, our studies suggest that CXADR expression on these two tumors increased survival times and that they act as ligands for the immune cells.

In contrast to the mouse, human tumor-infiltrating NK cells exhibit low levels of *AMICA1* across the seven tumors in the pan-cancer scRNA-seq dataset (Netskar et al., 2024). This discrepancy may reflect differences in the tumor microenvironment between mouse tumors and human tumors, particularly in the availability of cytokines capable of inducing *AMICA1* expression in NK cells. Furthermore, we see low JAML expression on NK cells within B16-F10 tumor and very high JAML expression on NK cells within the EO771 tumor suggesting that the tumor microenvironment can mediate JAML expression as both these tumors have undetectable levels of CXADR. Despite these differences, when we separate the JAML^+^ and JAML^-^ NK cells within the human pan-cancer NK cell dataset we see similar phenotypes that were seen in the mouse tumors, such as *CD27* and *PDCD1* are upregulated while *ITGAM* (encoding CD11b) is downregulated in JAML^+^ tumor infiltrating NK cells, suggesting that JAML^+^ NK cells are a small population of intermediate maturation of NK cells in humans. Other studies have seen higher expression of *AMICA1* in tissue resident NK cell subsets (Brownlie et al., 2021; Marquardt et al., 2019). Although we were unable to detect extracellular JAML with the only available antibody clone (401901) on human NK cells, we were able to detect intracellular JAML. These data suggest a specific signal, possibly a cytokine, that mediates the transport of JAML to the surface of human NK cells. In mice, we can detect extracellular JAML on splenic NK cells using IL-2, this same signal does not induce extracellular JAML on human blood NK cells, and a previous study suggests that IL-2 may downregulate or have no effect *AMICA1* transcript in human NK cells (Oesinghaus et al., 2025; Sabry et al., 2019). Thus, further research in the expression of JAML in human NK cells is needed to understand its transport to the surface and expression in response to the environment and cytokines.

Overall, the current study identifies JAML expression in both mouse and human NK cells within the tumor microenvironment, where it is shaped by ligand expression on the tumor cells, cytokine signaling in the NK cells, and inhibitory receptor engagement. similar to what has been seen in T cells, JAML expression is associated with checkpoint molecules such as PD-1 and LAG3, suggesting a link between JAML-CXADR interactions and checkpoint expression on the NK cells. Collectively, our findings support a model in which JAML-CXADR interactions play a role in NK cell anti-tumor activity and may be associated with the emergence of exhausted NK cells in the tumor microenvironment.

## Limitation of study

The current study demonstrates that mouse NK cells may express JAML when encountering certain tumors. It is unclear why some tumors induce JAML expression and others do not. It is clear however, that interaction of JAML with its ligand CXADR also reduces surface expression on NK cells which would skew interpretation of JAML data. Finally, while we can detect JAML RNA in intratumoral NK cells from humans, we do not see surface expression of JAML on NK cells. This also makes interpretation of the role of JAML on human NK cells difficult. It has been shown that JAML can engage multiple ligands, whereas our functional analyses focused exclusively on CXADR through genetic deletion, and therefore do not address potential contributions from other JAML-ligand interactions. However, the use of a JAML-Fc fusion protein to detect JAML ligands allows for unbiased recognition of all endogenous JAML-binding partners.

## Author Contributions

Conceptualization: D.L. A.K.W., C.T., J.C., M.J., K.K., B.J.C.; methodology: A.K.W., A.P. S.C., U.R., C.T., M.J., K.K., B.J.C; Data collection: A.K.W., S.S.A, N.M, J.B, D.M.B, A.P. L.N., J.F., L.K., A.H., C.B., R.M., R.L, C.P. S.P, Y.P. A.S.O, M.W., S.C., U.R. A.S.; Analysis and interpretation: D.L., A.K.W., S.S.A, N.M, J.B, D.M.B, L.N., J.F., L.K., A.H., C.B., R.M., U.R. S.C., A.S., M.J., B.J.C.; writing - original draft preparation, D.L., A.K.W., C.T. J.C. B.J.C.; critical revision of the article: A.K.W., K.K., J.C. B.J.C., Visualization, D.L., A.K.W., B.J.C; Funding acquisition, D.L., J.C. K.K. A.K.W. and B.J.C.; All authors have read and agreed to the published version of the manuscript.

## Key resources table

| REAGENT or RESOURCE | SOURCE | IDENTIFIER |
| --- | --- | --- |
| **Antibodies** |  |  |
| CD49a BV711 | BD Biosciences | Cat# 564863<br>RRID: 2738987 |
| Ly49f BUV395 | BD Biosciences | Cat# 745568<br>RRID: AB_2743089 |
| Ly49a Pacific Blue | Biolegend | Cat# 116810<br>RRID: AB_572013 |
| Ly49G2 PeCy7 | BD Biosciences | Cat# 560730<br>RRID: AB_1727551 |
| Ly49C | BD Biosciences | Cat# 748858<br>RRID: AB_2873261 |
| Ly49D BUV737 | BD Biosciences | Cat# 748496<br>RRID: AB_2872907 |
| Ly49H BV711 | BD Biosciences | Cat# 744264<br>RRID: AB_2742103 |
| Ly49I FITC | eBioscience | Cat# 11 5895 85<br>RRID: AB_465302 |
| CD69 FITC | BioLegend | Cat# 104505<br>RRID: AB_313108 |
| CD69 APC | BioLegend | Cat# 104514<br>RRID: AB_492843 |
| CD4 Per CP Cy 5.5 | BioLegend | Cat# 100434<br>RRID: AB_893324 |
| CD3e PE CF594 | BD Biosciences | Cat# 562286<br>RRID<br>AB_11153307 |
| CD45.2 AF700 | BioLegend | Cat# 109822<br>RRID: AB_493731 |
| TCR gamma/delta PE Cy5 | eBioscience | Cat# 15 5711 81<br>RRID AB_467570 |
| CD8a BUV615 | BD Biosciences | Cat# 613004<br>RRID: AB_2870272 |
| CD11c BUV395 | BD Biosciences | Cat# 564080<br>RRID: AB_2738580 |
| CD11b BV605 | BioLegend | Cat# 101257<br>RRID: AB_2565431 |
| CD317 (BST2) BUV805 | BD Biosciences | Cat# 749271<br>RRID: AB_2873648 |
| CD335 (NKp46) BV421 | BD Biosciences | Cat# 562850<br>RRID: AB_2737837 |
| NKp46 BV605 | BioLegend | Cat# 137619<br>RRID: AB_2562452 |
| CXCR6 PE Cy 7 | BioLegend | Cat# 151118<br>RRID: AB_2721669 |
| CX3CR1 BV650 | Biolegend | Cat# 149033<br>RRID: AB_2565999 |
| CD62L APC | BioLegend | Cat# 104412<br>RRID: AB_313099 |
| PD-1 BUV737 | BD Biosciences | Cat# 568607<br>RRID: AB_3684401 |
| CD25 BV510 | BioLegend | Cat# 102042<br>RRID: AB_2562270 |
| TIM3 BV711 | BD Biosciences | Cat# 747622<br>RRID: AB_2744188 |
| JAML PE | Miltenyi Biotec | Cat# 130-114-679<br>RRID: AB_2726744 |
| JAML APC | Miltenyi Biotec | Cat# 130-114-680<br>RRID: AB_2726745 |
| LAG3 SB645 | eBioscience | Cat# 64-2231-82<br>RRID: AB_2744872 |
| CD103 BV510 | BD Biosciences | Cat# 563087<br>RRID: AB_2721775 |
| Ly6C FITC | BD Biosciences | Cat# 553104<br>RRID: AB_394628 |
| GR1 Biotin | BioLegend | Cat# 108404<br>RRID: AB_313369 |
| 4-1BB PE Cy7 | eBioscience | Cat# 25-1371-82<br>RRID: AB_2573398 |
| CD39 APC | Miltenyi Biotec | Cat# 130-114-358<br>RRID: AB_2726584 |
| CD226 BV711 | BioLegend | Cat# 133609<br>RRID: AB_2715922 |
| I-Ab FITC | BD Biosciences | Cat# 553551<br>RRID: AB_394918 |
| KLRG1 PE Cy7 | BioLegend | Cat# 138416<br>RRID: AB_2561736 |
| NKG2A B6 APC | BioLegend | Cat# 142808 |
|  |  | RRID:<br>AB_11124538 |
| NK1.1 BV421 | BioLegend | Cat# 108732<br>RRID: AB_2562218 |
| TCF1/TCF7 BV421 | BD Biosciences | Cat# 566692<br>RRID: AB_2869822 |
| T-Bet RB545 | BD Biosciences | Cat# 569254<br>RRID: AB_3684909 |
| EOMES AF660 | Invitrogen | Cat# 606 4875 80<br>RRID: AB_2896280 |
| CD19 Biotin | eBioscience | Cat# 13-0191-85<br>RRID: AB_466385 |
| TER 119/erythroid Cells Biotin | BioLegend | Cat# 116204<br>RRID: AB_313705 |
| CD27 BUV661 | Invitrogen | Cat# 376-0271-82<br>RRID: AB_3074040 |
| CD127 BV650 | BioLegend | Cat# 135043<br>RRID: AB_2629681 |
| Streptavidin BV786 | BD Biosciences | Cat# 563858<br>RRID: AB_2869529 |
| Anti-CD16/CD32 | BD Biosciences | Cat# 553142<br>RRID: AB_394657 |
| Anti-CD3 FITC | BD Biosciences | Cat# 561806<br>RRID: AB_395739 |
| Anti-CD56 PE-Cy7 | BioLegend | Cat# 318318<br>RRID: AB_604099 |
| Anti-JAML BV421 | BD Biosciences | Cat# 749805<br>RRID: AB_2874059 |
| **Chemicals, peptides, and recombinant proteins** |  |  |
| JAML Fc-chimera | BioLegend | Cat#776806 |
| CXADR Fc-chimera | BioLegend | Cat#777602 |
| AffiniPure® F(ab')<sub>2</sub> Fragment Goat Anti-Human IgG PE | Jackson Human Research | Cat#109-116-048<br>RRID AB_2337676 |
| CD45 MicroBeads | Miltenyi Biotec | Cat#130-052-301 |
| rhIL2 | Biotechne | Cat#BT-002 |
| TAPI2 | Biotechne | Cat# 6013 |
| CFSE | ThermoFisher | Cat# C34554 |
| Cell trace Violet | ThermoFisher | Cat# C34557 |
| Live/Dead Fixable Aqua Dead Cell Stain | ThermoFisher | Cat# L34966 |
| Fixable Viability Dye eFluor 780 | ThermoFisher | Cat# 65-0865-14 |
| RPMI | HyClone | Cat# 16750-084 |
| FBS | Gibco | Cat# 10270-106 |
| 4% Formaldehyde | ITW Reagents | Cat# 252931.1211 |
| Intracellular staining | BioLegend | Cat# 421002 |
| Chromium-51 Radionuclide | Revvity | Cat#<br>NEZ030001MC |
| **Critical commercial assays** |  |  |
| NK cell isolation kit | Miltenyi Biotec | Cat# 130-115-818 |
| **Deposited data** |  |  |
| **Experimental models: Cell lines** |  |  |
| Mouse: MTAP1A | (Chambers, Grufman et al. 2007) |  |
| Mouse: MC57X | (Ciupitu et al. 2002) |  |
| Mouse: RMA-S | (Kärre et al., 1986) |  |
| Mouse:EO771 | ATCC | CRL-3461 |
| B16F10 | ATCC | CRL-6475 |
| YUMM1.7 | ATCC | CRL-3362 |
| **Experimental models: Organisms/strains** |  |  |
| Mouse: C57BL/6JRj | Janvier |  |
| Mouse: B6.*RAG1*<sup>-/-</sup> | Jackson Laboratory |  |
| Mouse: B6.b2m<sup>-/-</sup> | Jackson Laboratory |  |
| **Software and algorithms** |  |  |
| GraphPad Prism 9 | Graph Pad |  |
| FlowJo v10 | FlowJo, LLC |  |
| Biorender | Biorender | https://biorender.com |
| Seurat 3.0 | (Stuart, Butler et al. 2019) | https://satijalab.org/seurat/ |
| Swiss Model | (Waterhouse, Bertoni et al. 2018) | https://swissmodel.expasy.org/ |
| Scanpy 1.10.3 | (Wolf et al., 2018) | https://scanpy.readthedocs.io/en/latest/ |
| Celltypist 1.6.3 | (Xu et al., 2023) | https://www.celltypist.org/ |
| Python 3.10.13 | (Van Rossum and Drake, 2009) | https://www.python.org/ |
| Nextflow | (Di Tommaso et al., 2017) | https://www.nextflow.io/ |
| Nf-core RNA-seq | Patel *et al.* 2025 | https://zenodo.org/records/17909656 |
| DESeq2 | (Love et al., 2014) |  |
| Other |  |  |
| Protein Data base | https://www.rcsb.org | 3BIK, 4ZQK |
| CRISPOR | https://crispor.gi.ucsc.edu |  |

### Mice

C57BL/6 (B6, Janvier France), B6.*RAG1*^-/-^ (Mombaerts et al., 1992) and β2m-/- (Koller et al., 1990) were housed under specific pathogen free conditions at the Astrid Fagraeus Laboratories, Karolinska Institutet, Stockholm. All procedures were performed under both institutional and national guidelines (Ethical numbers from Stockholm County Council). Sex and aged match mice were used for all experiments. Mice were chosen randomly for control or treated groups.

### Human subjects and processing of blood

Human peripheral blood mononuclear cells (PBMCs) were isolated by density gradient centrifugation using Lymphoprep (Stemcell Technologies). Ethical permission to study PBMCs from healthy donors was obtained from the regional ethical committee in Stockholm (2020-02604). PBMCs used in this study came from Karolinska Universitetssjukhuset.

### Tumors

MHC-I-deficient lymphomas RMA-S (*TAP2*-deficient)(Kärre et al., 1986), and *TAP1*-deficient MCA fibrosarcoma (clone MTAP1A)(Chambers et al., 2007), MCA fibrosarcoma (clone MC57X)(Wen et al., 1997), EO771 (Johnstone et al., 2015), YUMM1.7(Meeth et al., 2016) and B16F10 have all been described previously. Since the sex of the tumors was unknown, male mice were used as recipients except for EO771 where female mice were used. Mice were given RMA-S, MTAP1A, YUMM1.7, and MC57X at 10^5^ and B16F10 at 5×10^5^ tumors s.c. and EO771 in the mammary fat pad at 2×10^5^. Tumor growth was measured every two days and mice were sacrificed when the tumor reached 10^3^ mm.

### NK cell purification and culture

Single-cell suspension from spleens was depleted of erythrocytes, and NK cells were positively sorted by negative sorting using MACS separation, according to the manufacturer’s instructions (Miltenyi Biotec, Bergisch Gladbach, Germany). Cells were resuspended in complete medium (MEMalpha; 10 mM HEPES, 2 × 10^−5^ M 2-ME, 10% FCS, 100 U/ml penicillin, 100 U/ml streptomycin) with 1000U/ml IL-2 (RNDsystems). In some experiments, 100 ng/ml mouse IL-12 (PeproTech), and 100 ng mouse IL-21 (PeproTech) were added for the final two days. For isolation of NK cell subsets, NK cells were isolated as above and then sorted on BD Influx (Becton Dickenson, CA, USA).

### Flow Cytometry and Imaging Flow Cytometry

Splenic and tumor NK cells were stained aÅer single cell prepara∼ons were depleted of erythrocytes. Cells were stained as outlined in the various experiments and the an∼bodies used outlined above. For cultured NK cells, cells were first collected at the ∼mes outlined above and then stained. Flow cytometry was performed on CyAN ADP LX 9-colour flow cytometer (Beckman Coulter, Pasadena, CA), BD FACSCanto (BD Biosciences) or Sony 7000 Spectral Cell Analyzer (Sony Biotechnology San Jose CA). Data were analyzed using FlowJo v10 soÅware (BD Biosciences). Imaging flow cytometry was performed on an Amnis Imagestream Mk II (Cytek) and the data was analyzed using IDEAS soÅware (Cytek).

### *Cxadr* KO in MC57X and MTAP1A mouse tumours

*Cxadr* KO MC57X and MTAP1A knock-out (KO) cells were generated by electroporating the cells with CRISPR/Cas9 components as a ribonucleoprotein (RNP) using a Lonza 4D-Nucleofector system (4D-Nucleofector Core Unit: #AAF-1002B; 4D-Nucleofector X Unit: #AAF-1002X, Lonza Group AG). Exon 1 of *Cxadr* was targeted with a gRNA (5’ GAAGCACAGTAGGCGCGCCA) designed with the help of CRISPOR tool (Concordet and Haeussler, 2018). Cas9 protein containing three nuclear localisation signals was produced in-house by the Protein Science Core Facility^2^. Alt-R^TM^ tracrRNA (#1072533, Integrated DNA Technologies) and Alt-R^TM^ crRNA (IDT) were annealed 1:1 ratio at 95 °C for 5 minutes to form 100 µM gRNA. Cas9 and gRNA were incubated for 15 minutes at RT in a ratio of 1:1.2, after which the RNP was nucleofected into MC57X or MTAP1A cells resuspended in Gene-Pulser Electroporation buffer (#165-2676, Bio-Rad Laboratories, Inc.) using pulse code CM137. KO efficiency was assessed by flow cytometry on day seven and CXADR-negative cells were sorted at the Biomedicum Flow cytometry Core facility using the JAML-Fc protein (Biolegend)

### mRNA library construction

Total RNAs from enriched NK cells were extracted using Direct-zol RNA miniprep kit (Zymo Research) following the manufacturer’s instructions. The quality of the input RNAs were then assessed on Bioanalyzer using RNA 6000 nano kit (Agilent Technologies). Total RNAs were quantified using Qubit high sensitivity RNA Assay kit (Thermofisher Scientific). 100 ng total RNA was then used to construct single-indexed mRNA library using TruSeq Stranded mRNA kit (Illumina). The protocol was carried out following the standard manufacturer’s instruction. The quality and the median library fragment size were assessed on Bioanalyzer using High Sensitivity DNA analysis Kit (Agilent technologies). Quantitation of libraries was performed by quantitative PCR using Collibri Library Quantification Kit (Invitrogen).

All libraries were then pooled at equal concentration with 2% Phi-X spiked into the library pool and subsequently loaded on an Illumina Nextseq 550 platform for paired-end sequencing with read length of 75 cycles from each end.

### Cytotoxicity assay

Tumor cells were incubated for 1 h in the presence of Na_2_^51^CrO_4_ (Revvity, Waltham, MA) and then washed thoroughly in cell culture media. The tumor cells were added to NK cells at the ratios indicated in a U-bottom plate and the plate centrifuged for 300g for 1 min. After 3h, supernatants these wells were collected and analyzed in triplicate by using a gamma radiation counter (Wallac Oy, Turku, Finland). Specific lysis was calculated according to the following formula: % specific lysis=(experimental release-spontaneous release)/(maximum release-spontaneous release)x100.

### scRNA-seq and RNA-seq analyses

Data for the scRNA-seq pan-cancer dataset were retrieved in h5ad format from https://doi.org/10.5281/zenodo.8434223. The scRNA-seq data was analyzed using Scanpy 3.10.3 and the different tissues were subset based on the metadata provided in the h5ad file. The NK cells were from tumor or healthy tissue were further analyzed to identify clusters using Leiden algorithm and identify clusters enriched in *JAML* expression. Pseudobulk differential gene expression of JAML- and JAML+ tumor infiltrating NK cells was prepared by subsetting the NK cells based on a JAML expression of <0.1 for JAML^lo/-^ and >=0.1 for JAML^+^ and then those NK cell groups were randomly separated into 3 groups to create pseudoreplicates and the python package of DESeq2 was used to determine differentially expressed genes between the NK cells. Data for the RNA-seq data were retrieved from project SRP125125 and then count matrices were generated from the fastq files using the nextflow RNA-seq pipeline and hg38 genome build. In short, adapter trimming and QC was performed on the fastq files and then the reads were aligned using STAR and the was specific isoform *JAML* transcripts were quantified using salmon and displayed as transcript per million (TPM) expression.

### Statistics

All statistical analysis was performed using GraphPad Prism software (La Jolla, CA)

## Code availability

The code generated for our analyses for the IL-2 stimulated NK cell RNA-seq dataset and pan-cancer scRNA-seq dataset is available on GitHub at X.

## Data availability

The RNA-seq data generated for this study have been deposited in the GEO under accession code GSEXXXX. Source data of the DEGs from the IL-2 stimulated NK Cells DEG are provided with this paper.

## Acknowledgements

This work was funded by the Swedish Cancer Society (B.J.C., J.C., K.K.) and Swedish Research Council (J.C.).

Daniel Labuz is postdoctoral position sponsored by Swedish Cancer Society.

Carmela Passacatini was supported by SG-2021-1237555/Ministero della Salute; Ricerca corrente/Ministero della Salute (Ministry of Health, Italy).

The authors would like to acknowledge the contributions of the Biomedicum Flow cytometry Core facility, financed by the Infrastructure Board at Karolinska Institutet, for providing access to equipment, cell sorting service and technical expertise.

Part of this work was facilitated by the Protein Science Facility at Karolinska Institutet, Stockholm, (and we would like to thank Dr Emilia Strandback, Dr Henry Ampah-Korsah, Dr Henrik Spåhr and Dr Tomas Nyman for assistance).

## Notes

Financial Support: This work was funded by the Swedish Cancer Society, Swedish Research Council, the Karolinska Institute Foundations

### Competing Interest Statement

The authors have declared no competing interest.

