## Supplemental Figures 1-5 for "Intratumoral expression of JAML on NK cells is controlled by tumor microenvironment and MHC class I interaction"

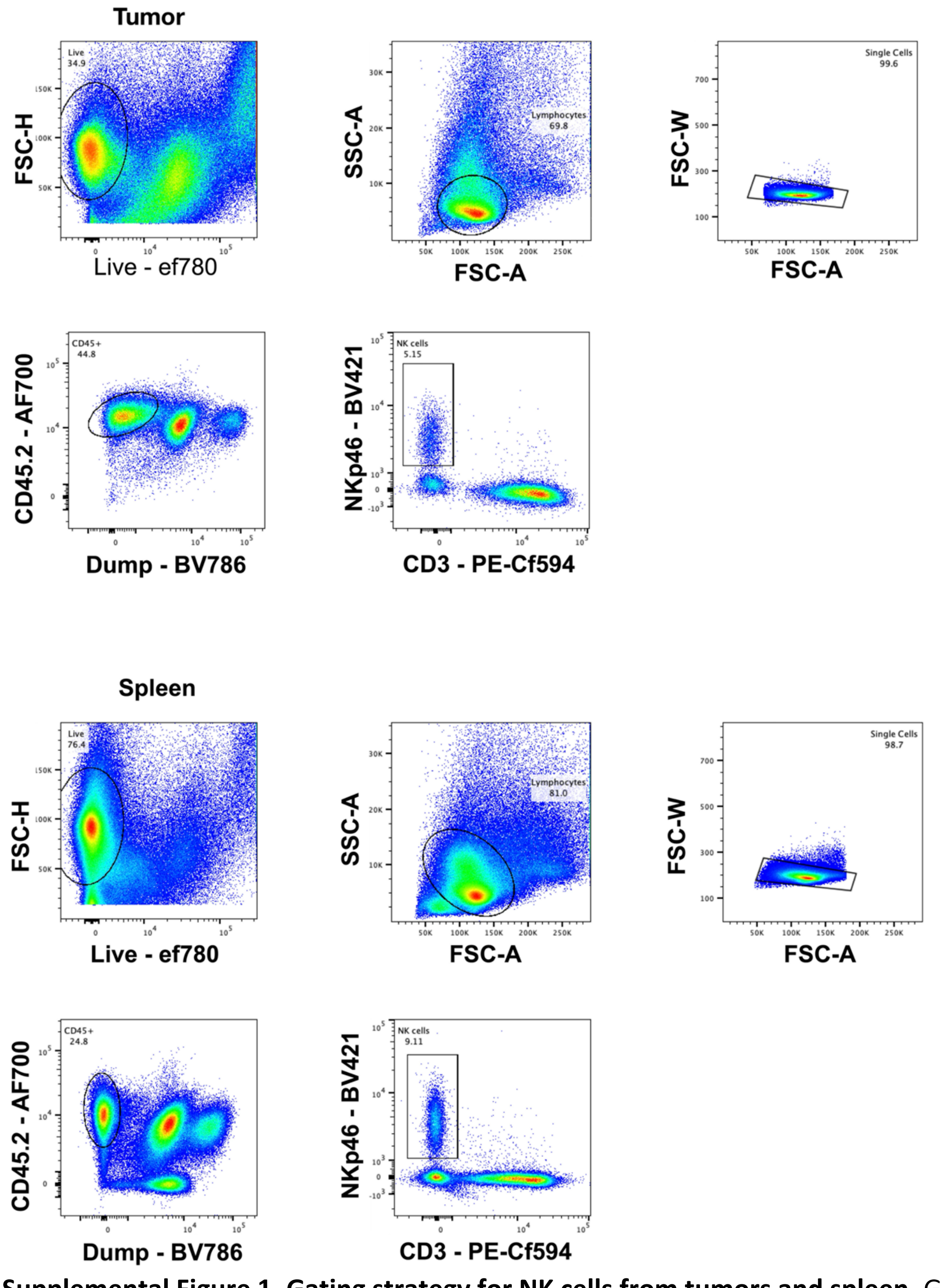


**Supplemental Figure 1. Gating strategy for NK cells from tumors and spleen.** Gating was performed on live cells and then on size. Doublets were then removed by gating on the forward scatter area and width. B cells, granulocytes and red blood cells were gated out using anti-CD19, GR1, and Ter119 respectively. NK cells were then selected as being NKp46$^+$CD3$^-$.

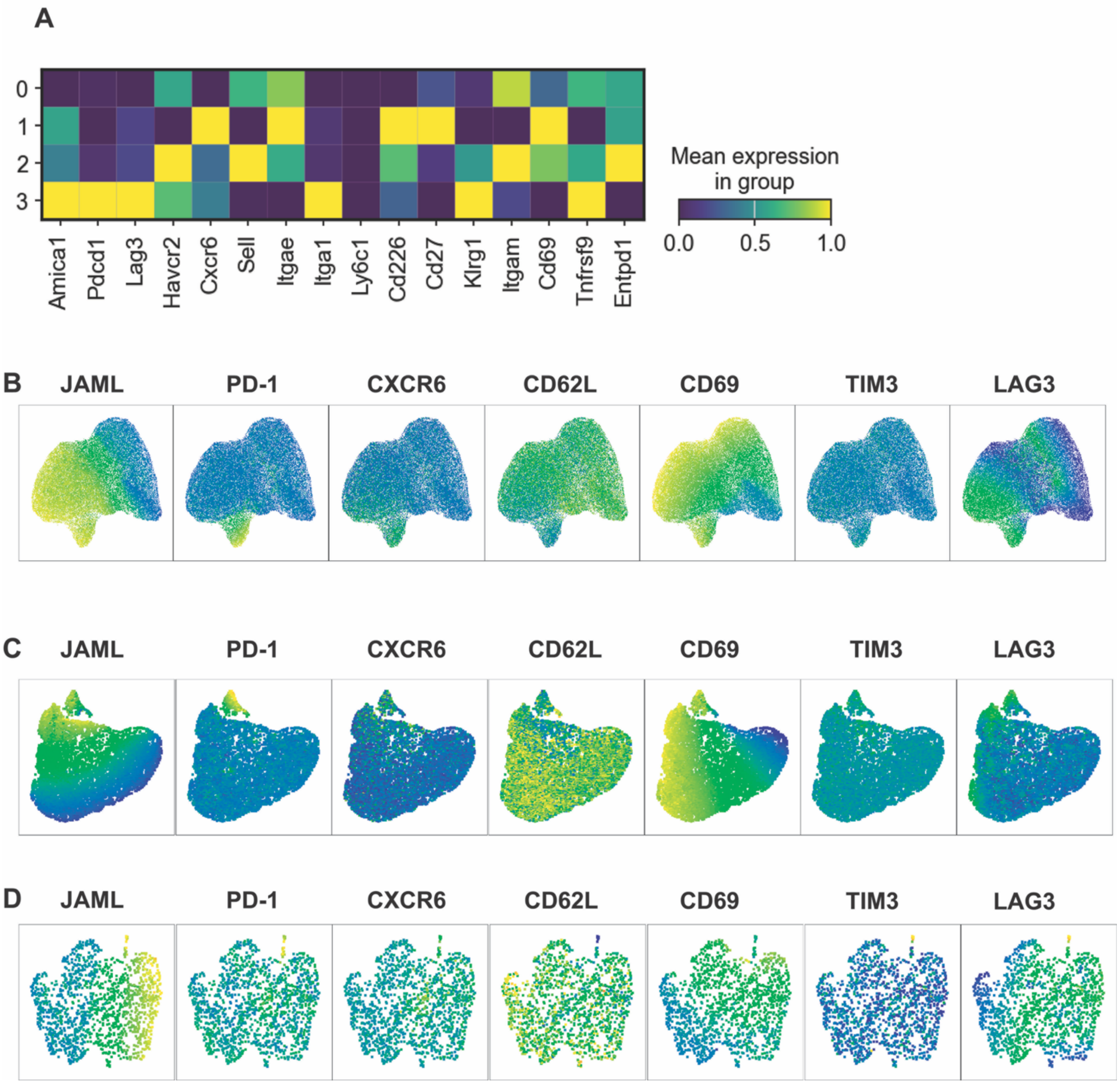


**Supplemental Figure 2. Phenotype of JAML expressing NK cells.** (A) Matrix plot of scRNAseq of intratumoral NK cells from EO771 tumors. (B) UMAP of NK cell populations examining the distribution of JAML, PD-1, CXCR6, CD69 CD62L, TIM3, Lag3 by flow cytometry from concatenated NK cells from EO771 (n=6), (C) MC57X (n=5) and (D) YUMM1.7 (n=3)

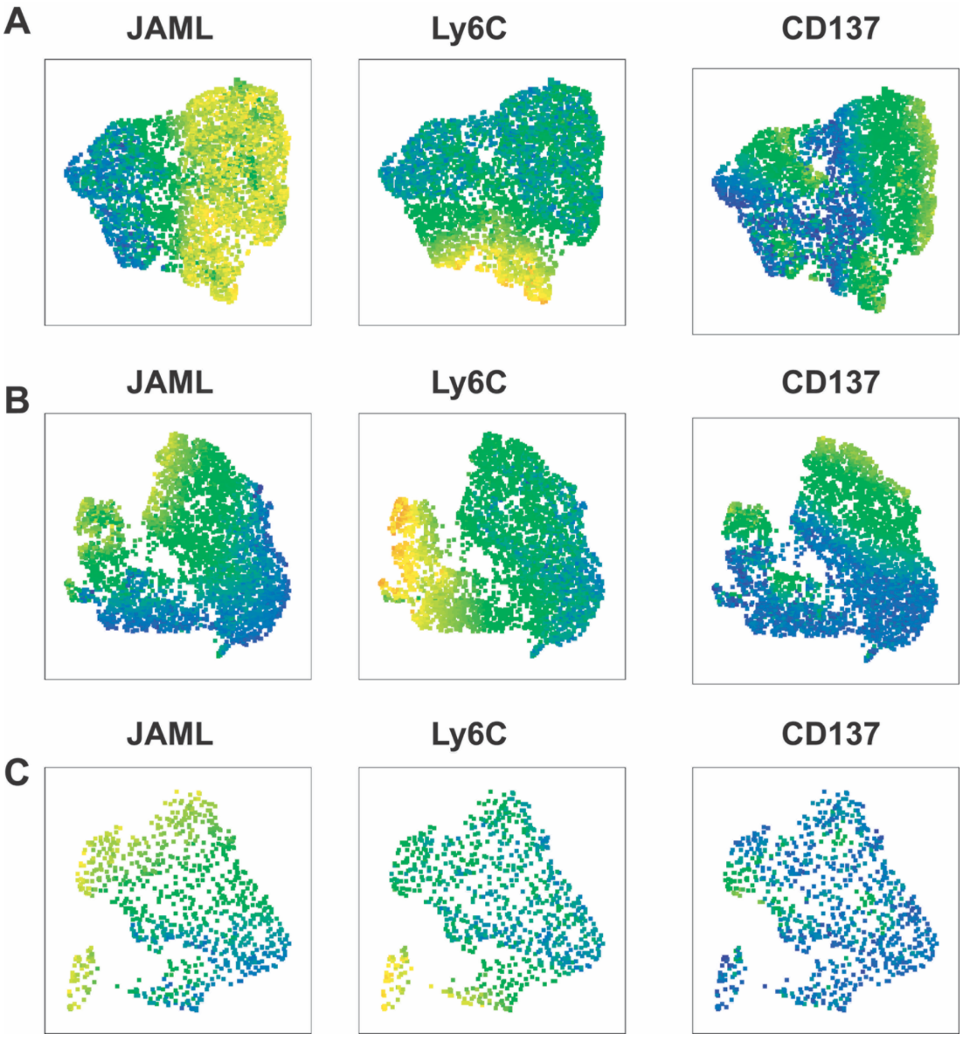


**Supplemental Figure 3. Phenotype of JAML expressing NK cells.** (A) UMAP of NK cell populations examining the distribution of JAML, Ly6C, and CD137 by flow cytometry from concatenated NK cells from EO771 (n=6), (B) MC57X (n=5) and (C) YUMM1.7 (n=3)

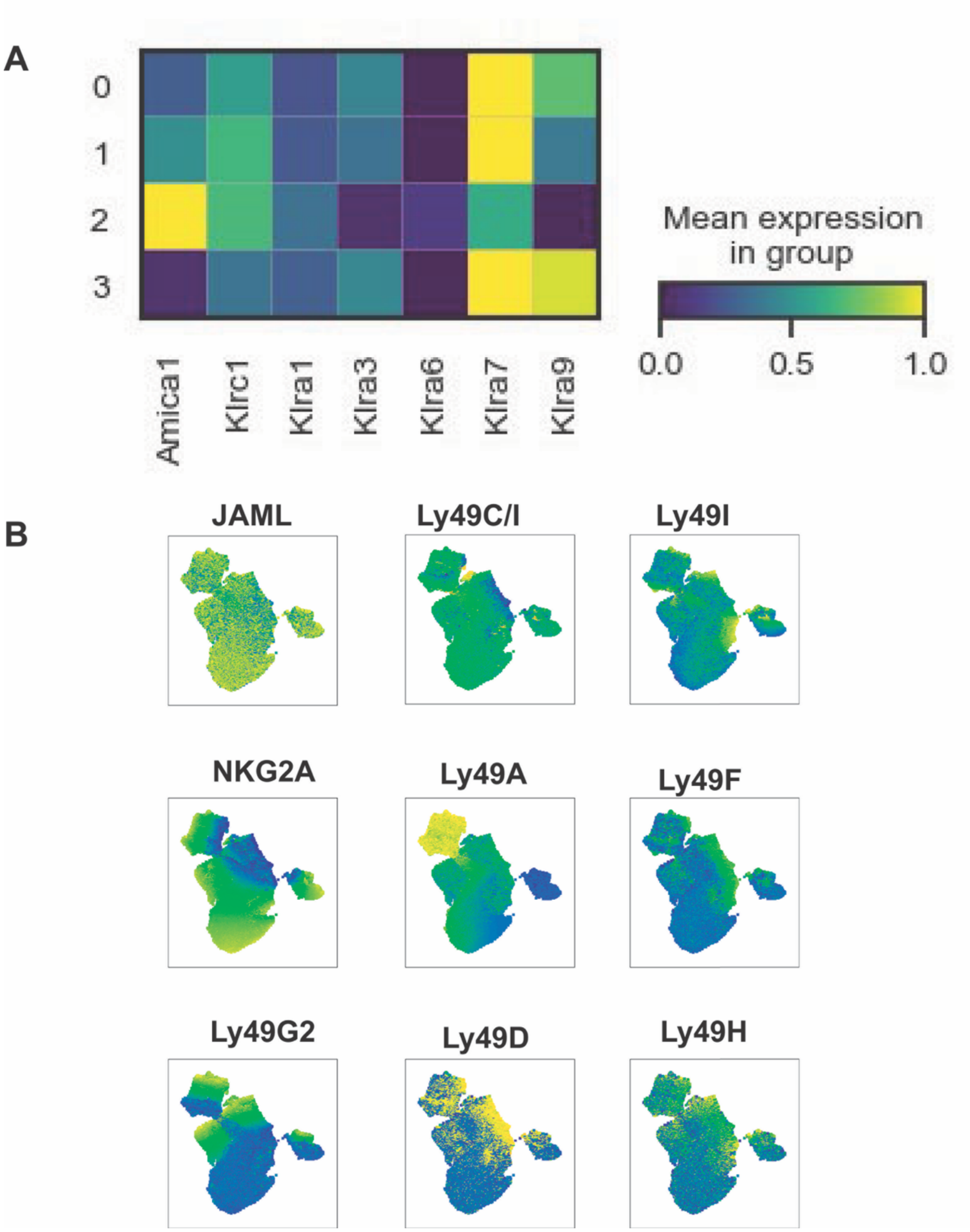


**Figure 4. JAML expression on Ly49 expression of the NK cells from EO771.** (A) Matrix plot of expression of cytokine receptors and JAML (*AMICA1*) on NK cells from EO771. (B) UMAP plots of JAML, the inhibitory Ly49s Ly49C/I, Ly49I, NKG2A, Ly49A, Ly49F, Ly49G2 and the activating Ly49s Ly49D and H. UMAP was performed using the JAML and the negative Ly49s (n=6).

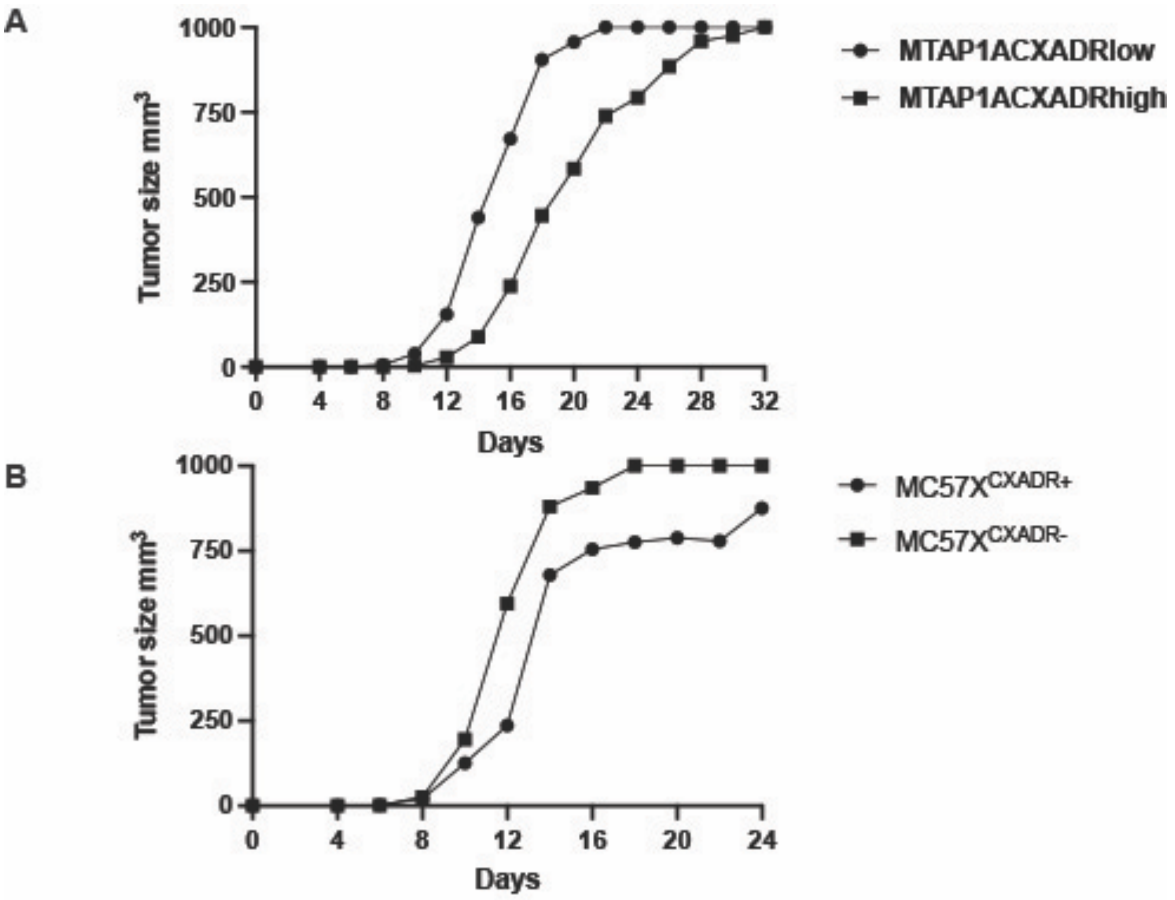


**Figure 5. Lack of CXADR on tumors leads to faster tumor outgrowth.** Growth curves of (A) MTAP1A$^{CXADRlow}$ and MTAP1A$^{CXADRhigh}$ tumors and (B) of MC57X$^{CXADR-}$ and MC57X$^{CXADR+}$

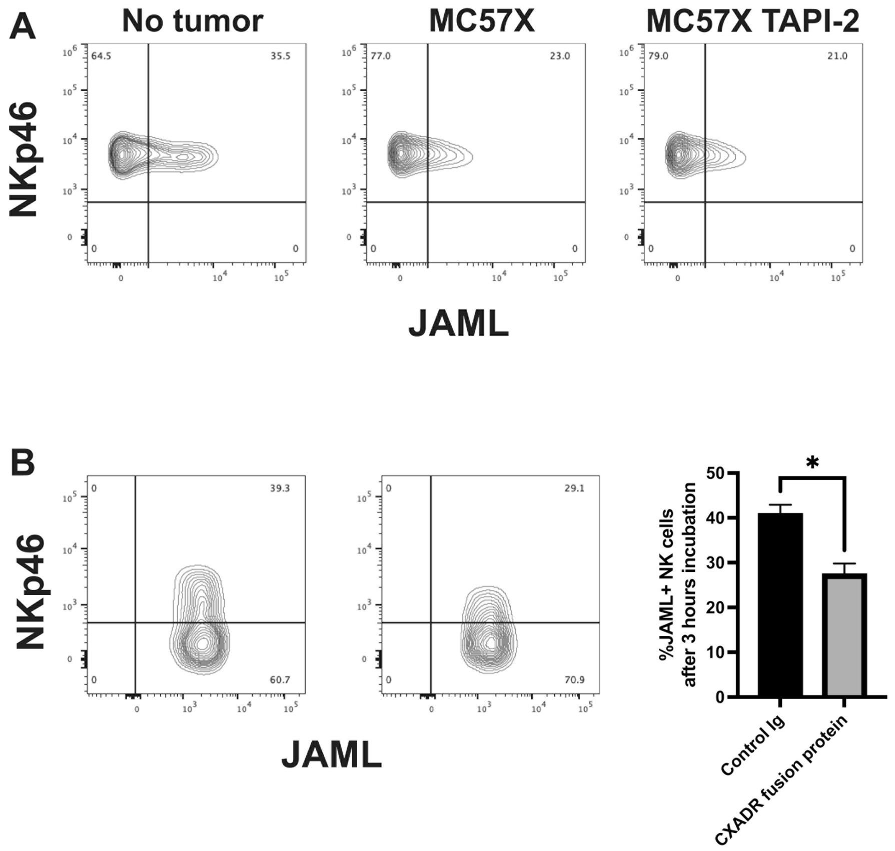


**Figure 6. JAML expression is reduced by interaction with bound CXADR but not metalloprotease inhibitor TAPI-2.** (A) JAML expression of JAML in the presence or absence of TAPI-2.B) JAML expression on NK cells exposed to place bound CXARDR-fusion protein or control Ig (*p<0.05 paired t-test).
